# Phase transition of spindle pole localized protein orchestrates nuclear organization at mitotic exit

**DOI:** 10.1101/2025.01.22.634232

**Authors:** Ashwathi Rajeevan, Vignesh Olakkal, Madhumitha Balakrishnan, Dwaipayan Chakrabarty, François Charon, Daan Noordermeer, Sachin Kotak

## Abstract

Animal cells dismantle their nuclear envelope (NE) at the beginning and reconstruct it at the end of mitosis. This process is closely coordinated with spindle pole organization: poles enlarge at mitotic onset and reduce size as mitosis concludes. The significance of this coordination remains unknown. Here, we demonstrate that Aurora A maintains a pole-localized protein NuMA in a dynamic state during anaphase. Without Aurora A, NuMA shifts from a dynamic to a solid phase, abnormally accumulating at the poles, leading to chromosome bending and misshaped nuclei formation around poles. NuMA localization relies on interactions with dynein/dynactin, its coiled-coil domain, and intrinsically disordered region (IDR). Mutagenesis experiments revealed that cation-π interactions within IDR are key for NuMA localization, while glutamine residues trigger its solid-state transition upon Aurora A inhibition. This study emphasizes the role of the physical properties of spindle poles in organizing the nucleus and genome post-mitosis.

## Introduction

Eukaryotic cells maintain and protect their genetic information within a single nucleus for most of the cell cycle. Within the nucleus, chromosomes adopt a complex 3D organization (Cremer and Cremer, 2010), and changes in nuclear shape can influence gene regulation (Akhtar and Gasser, 2007; Almonacid et al., 2019, Tajik et al., 2016). At mitotic onset, animal cells condense their genome and disassemble the nuclear envelope (NE) (Antonin et al., 2016; Paulson et al., 2021). Concurrently, spindle poles (referred to as “poles”) are reinforced to produce robust microtubule asters, ensuring that all chromosomes are efficiently and rapidly captured (Woodruff et al., 2014; Conduit et al., 2015). During mitotic exit, the microtubule nucleation capacity of the poles diminishes, and the nuclear envelope reforms. This coordination between NE disassembly/reassembly and spindle pole organization is essential for successful mitosis; however, the consequences of failed coordination are unclear.

Spindle organization during mitosis is governed by the pole-localized protein NuMA (Merdes et al., 1996; Merdes et al., 2000; Hueshen et al., 2017; Kiyomitsu and Boerner, 2021). NuMA levels progressively increase at the poles during mitotic entry and decrease at mitotic exit (Kotak et al., 2013). Recent research indicates that NuMA’s accumulation at the poles may be influenced by its ability to undergo liquid-liquid phase separation (LLPS) *in vitro*, driven by its intrinsically disordered C-terminal region (Sun et al., 2021; Ma et al., 2022). However, it remains uncertain whether the physiological concentration of NuMA (<1 µM; Hein et al., 2015) is sufficient to drive its accumulation at the poles solely via LLPS. A key mitotic kinase, Aurora A, is crucial in maintaining NuMA levels at the poles during metaphase. In the absence of Aurora A, NuMA abnormally accumulates at the poles (Gallini et al., 2016; Kotak et al., 2016). However, it remains unknown whether Aurora A activity is continuously required to regulate NuMA pole accumulation after mitosis. Importantly, the potential consequences of failing to dissolve poles located near newly forming nuclei during mitotic exit have not been explored in any cellular system.

## Results

### Aurora A activity during anaphase is essential for proper nuclear organization

While analyzing Aurora A role in mitosis, we noted that a majority (95%) of HeLa cells that rapidly exit mitosis (within 45-60 min) in the presence of a specific Aurora A inhibitor MLN8237 (50 nM) harbored abnormal organization of the newly formed nuclei without showing any sign of chromosome instability and chromosome missegregation errors (Fig. 1 A-D). This result indicates that Aurora A activity during anaphase before nuclear envelope reformation (NER) is crucial for proper nuclear organization. Supporting this notion, we detected significant levels of active auto- phosphorylated Aurora A at the poles in anaphase (Fig. S1 A, B). To explore the role of Aurora A during anaphase before NER, we established a novel genetic tool to acutely remove the protein selectively during anaphase. Building on previous studies of sea urchin CyclinB1, where the N-terminus was shown to be sufficient for proteasome-mediated degradation during anaphase (Glotzer et al., 1991), we identified the first 86 amino acids of human CyclinB1 (referred to as CycB) as containing the probable degron sequence. This was validated by fusing this region to AcGFP (referred to as CycB-AcGFP), which led to the rapid degradation of CycB-AcGFP during anaphase (Fig. 1 E, F and Fig. S1 C, D). Next, we fused Aurora A (67-403 amino acids) to CycB and AcGFP (referred to as CycB-Aurora A^r^-AcGFP; Fig. 1E) and generated transgenic cell lines. The N-terminal sequence (1-66 amino acids) of Aurora A was intentionally removed because (i) it contains a recognition site for the APC/C^Cdh1^ (Littlepage and Ruderman, 2002), and we intended to drive Aurora A degradation solely by CycB E3 ubiquitin ligase APC/C^Cdc20^, (ii) a monoclonal antibody against the N-terminus of Aurora A can be used to detect endogenous Aurora A, and (iii) an siRNA against the N-terminus of Aurora A can deplete endogenous protein. We chose an engineered cell line that expresses catalytically active CycB-Aurora A^r^-AcGFP in amounts indistinguishable from the endogenous Aurora A (Fig. 1G) and localizes similarly to endogenous Aurora A (Fig. 1F; Fig. S1E). CycB-Aurora A^r^-AcGFP significantly rescued the mitotic index and chromosome instability errors seen upon endogenous protein depletion (Fig. S1 F, G). As expected, CycB-Aurora A^r^-AcGFP is swiftly degraded during the metaphase-to-anaphase transition (Fig. 1 F, H). Cells expressing CycB-Aurora A^r^-AcGFP failed to accurately separate their chromosomes (Fig. S1H), a phenotype linked to Aurora A function in anaphase (Reboutier et al., 2011). These experiments established that CycB-Aurora A^r^-AcGFP expressing cells can be utilized to study Aurora A-dependent function/s at mitotic exit. Intriguingly, a significant number of CycB-Aurora A^r^-AcGFP expressing cells did not show any chromosome instability errors but contained misshaped nuclei (Fig. 1 I-K). This confirms that Aurora A activity in anaphase is essential for proper nuclear organization.

**Fig. 1.**
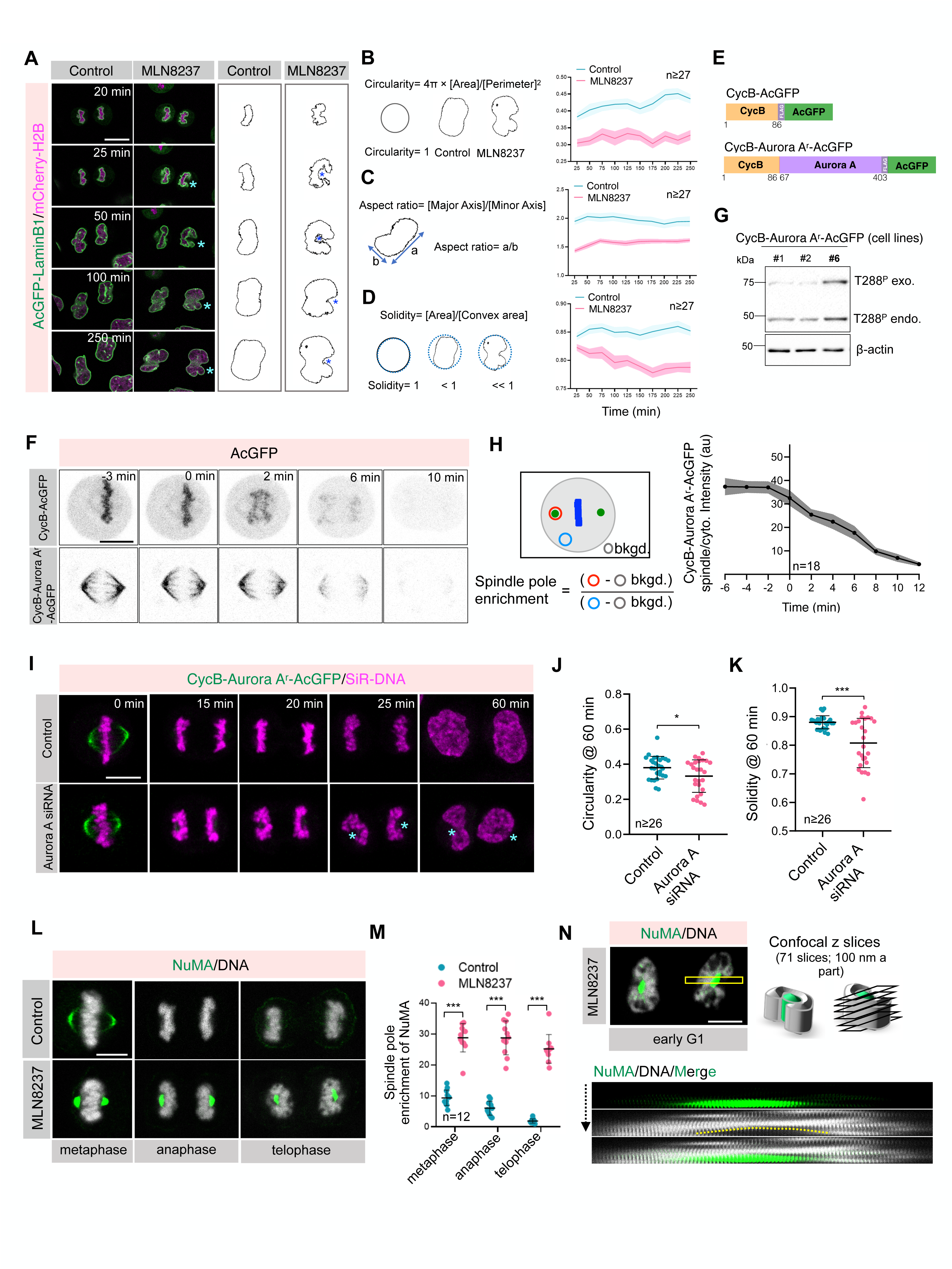
Aurora A activity is vital for proper nuclear morphology at the mitotic exit (**A**) Panels from time-lapse movies of representative DMSO (control) and MLN8237-treated HeLa cells stably coexpressing AcGFP-tagged LaminB1 (green) and mCherry-H2B (magenta). In this and other figure panels, timepoint t=0 min was set to the metaphase-to-anaphase transition. Asterisks in MLN8237-treated cells depict the bending of chromosome ensemble mass around the position where poles are usually located. The shape of the chromosomes ensemble is shown on the right in these conditions. (**B-D**) Morphological analysis of various nuclear parameter [circularity (B), aspect ratio (C), and solidity (D)] in control and MLN8237-treated cells at different time points post metaphase-to-anaphase transition. Curves and shaded areas indicate mean ± SEM. *P* values for circularity and aspect ratio for all the time points is <0.001, however, the *P* values for solidity at 25 min post metaphase to anaphase is =0.028, and for the rest of the time points <0.001. (E) Schematic representation of CycB and siRNA resistant form of Aurora A construct with AcGFP- and mono FLAG-tag at the C-terminus. (F) Confocal live-cell imaging of HeLa cells stably expressing CycB-AcGFP or CycB-AuroraA^r^-AcGFP. (G) Immunoblot analysis of mitotic cell extracts of HeLa cells stably expressing three independent clones (#1, #2, and #6) of CycB-Aurora A^r^-AcGFP. Extracts were probed with antibodies directed against autocatalytically active Aurora A (T288^P^) and b-actin. Endogenous and exogenous T288^P^ bands are indicated. Line #6 is utilized for all future analyses. The molecular mass is indicated in kilodaltons (kDa) on the left. (H) Schematic representation of the method for quantifying spindle pole intensity of CycB-Aurora A^r^-AcGFP [in arbitrary unit (au)] and the outcome of such analysis over time. Curve and shaded areas indicate mean ± SEM. bkgd., representing background intensity. (I) Confocal live-cell imaging of HeLa cells stably expressing CycB-AuroraA^r^-AcGFP (green) and probed for silicon-rhodamine DNA (SiR-DNA; magenta) to visualize chromosomes ensemble during anaphase in control and upon transfection with Aurora A siRNA for 60 h. (**J, K**) Nuclear shape analysis [circularity (J) and solidity (K)] from the confocal live-cell imaging of cells mentioned in panel I. Error bar indicates SD. In this and other Figures, ns- p>0.05; *-p<0.05; **-p<0.01; ***- p<0.001 as determined by two-tailed Student’s t-test. (**L, M**) Immunofluorescence (IF) analysis of different cell cycle stages of HeLa cells stained for NuMA (green) and DNA (grey) in control and in MLN8237 (50 nM) (L). Spindle pole intensity of NuMA (M; in au) was calculated at these different stages as shown for Fig. 1H. Error bar indicates SD. (**N**) IF analysis of HeLa cell stained for NuMA (green) and DNA (grey) in the early G1 phase of the cell cycle upon MLN8237 treatment. The midplane section of the cell is shown. The schematic on the right represents an area (marked by yellow in the image) covered from top to bottom and sliced into 71 z-projections with a step size of 100 nm to build the kymograph on the bottom. See also figs. S1-S3. Scale bars 10 *μ*m.

### Aurora A dissolves pole-localized NuMA in anaphase to ensure the proper nuclear organization

We noticed that MLN8237-treated and CycB-Aurora A^r^-AcGFP expressing cells showed peculiar bending of segregated chromosomal mass at the position where generally poles are present (Fig. 1 A, I-indicated by asterisks). Earlier, Aurora A inhibition in metaphase was shown to abnormally increase NuMA levels at poles (Gallini et al., 2016; Kotak et al., 2016). Therefore, we characterized pole-localized NuMA and the segregated mass of chromosomes ensemble in MLN8237-treated HeLa cells at anaphase onset. NuMA, which usually swiftly dissolves at poles, is significantly enriched at the poles upon Aurora A inhibition in anaphase (Fig. 1 L, M). Notably, the segregated chromosome sets had bent around the abnormally localized NuMA at telophase and early G1 phase (Fig. 1 L, N). A similar observation was made in several other cell lines (Fig. S2 A-D). Geometric measurements showed that NuMA geometry at the poles significantly differs in Aurora A inhibited cells (Fig. S2 E, F). This abnormal accumulation of NuMA at poles is not because of excess NuMA in MLN8237-treated cells (Fig. S2G) but due to overall decreased levels of NuMA in the cytoplasm (Fig. S2H).

To directly visualize the impact of NuMA pole-accumulated NuMA on nuclear morphology in Aurora A inhibited cells, we imaged NuMA and AcGFP-LaminB1 by live-cell imaging. In control cells, AcGFP-LaminB1 localized around the segregated mitotic chromosome ensemble at ∼15 min post anaphase onset; during this time, NuMA levels at the poles were significantly diminished. In contrast, in Aurora A inhibited cells, NuMA at the poles was readily visible for a considerably longer duration of ∼1 h or more, and the newly formed nuclei were bent around the NuMA-localized poles (Fig. S2 I, J). Similar observations were made in cells coexpressing mCherry-NuMA with nuclear envelope marker GFP-Nup107 (Fig. S3 A, B). Moreover, to link NuMA pole accumulation with Aurora A function during anaphase, we generated stable cells coexpressing CycB-Aurora A^r^-AcGFP and mCherry-NuMA. Consistent with the above, NuMA levels were significantly enriched at the pole in cells expressing CycB- Aurora Ar-AcGFP upon endogenous Aurora A depletion (Fig. S3 C, D).

Aurora A phosphorylates NuMA at Serine 1969 in its C-terminus (Gallini et al., 2016; Kettenbach et al., 2011). Therefore, we tested whether the expression of NuMA containing the S1969A replacement with AcGFP (AcGFP-NuMA^r^S1969A) would lead to its accumulation at the poles during anaphase, similar to anaphase Aurora A inhibition. If so, could the accumulation of AcGFP-NuMA^r^S1969A at the poles bend the segregated chromosome sets? As envisaged, AcGFP- NuMA^r^S1969A robustly accumulated at the poles in contrast to wild-type NuMA, and pole-localized AcGFP-NuMA^r^S1969A could bend the segregated chromosome sets and the nascent nuclei (Fig. S3 E, F). Based on these results, we conclude that persistent Aurora A activity in anaphase dissolves pole-localized NuMA. Otherwise, NuMA abnormally accumulates behind the segregating chromosome sets, creating a barrier for the newly formed nuclei to attain proper morphology.

### Aurora A regulates the phase transition of NuMA at poles

Transiently expressed GFP-NuMA shows significantly slower recovery at the poles in fluorescence recovery after photobleaching (FRAP) experiments upon Aurora A inhibition during metaphase (Gallini et al., 2016). We obtained similar results in cells stably expressing AcGFP-tagged NuMA in cells acutely treated with MLN8237 in metaphase and during anaphase (Sana et al., 2022) (Fig. 2 A-C and Fig. S4 A-D). Next, we performed a ‘half-bleach’ experiment (Brangwynne et al., 2009) by bleaching AcGFP-NuMA within the pole and then analyzing fluorescence recovery within the photo-manipulated area over time (Fig. S4 E, F). The Aurora A-inhibited cells were significantly impaired in regaining the AcGFP-NuMA fluorescence signal within the poles (Fig. S4 G, H). Furthermore, cells expressing NuMA with photoconvertible fluorescent protein mEOS failed to acquire photoconverted red fluorescence from the non-irradiated pole in Aurora A inhibited cells (Zhang et al., 2012) (Fig. 2 D, E).

**Fig. 2.**
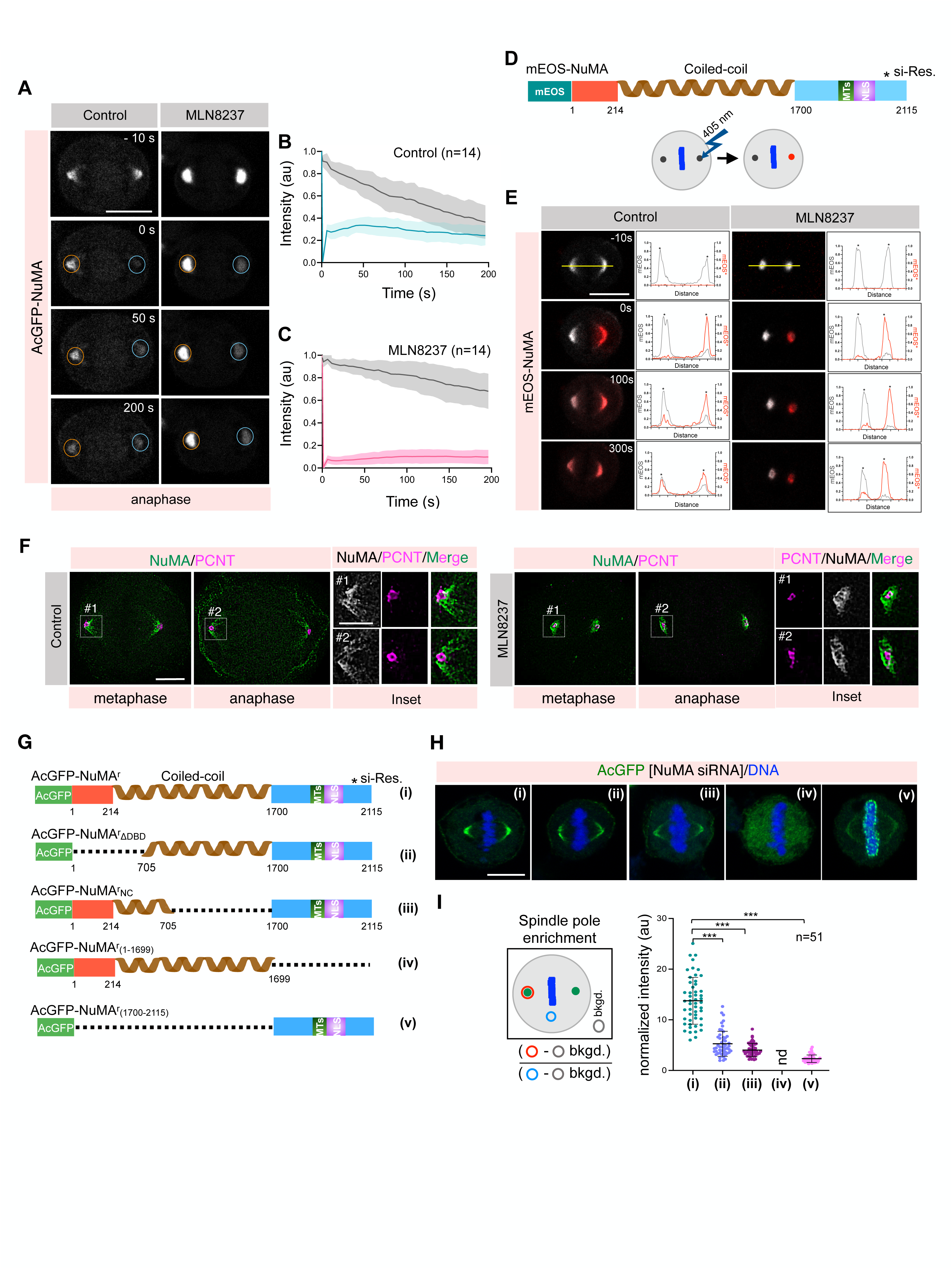
**Aurora A regulates the phase transition of NuMA at the poles** (**A-C**) FRAP analysis of HeLa cells stably expressing AcGFP-NuMA (grey) in control and upon Aurora A inhibition with MLN8237 during anaphase. Time is indicated in seconds (s). The unbleached and bleached regions of the cell are shown by orange and blue circles, respectively. The AcGFP recovery profile of the bleached area for control (B) and MLN8237-treated cells (C) is shown on the right. Curves and shaded areas indicate mean ± SD. (**D, E**) Domain organization of mEOS-tagged NuMA (mEOS-NuMA). Microtubule interaction and nuclear localization signals are shown as MTs and NLS, respectively. As shown in the schematics, mEOS-NuMA was photoconverted from green (shown as grey) to red at one pole by the exposure of 405 nm laser (D), followed by assessing the accumulation of photoconverted mEOS-NuMA on the non-photoconverted pole by confocal live-cell imaging (E). Line-scan analysis of the poles before and after photoconversion is shown for the unperturbed and perturbed poles in untreated and MLN8237-treated cells. More than 30 cells were analyzed; the representative cell is shown here. (F) Super-resolution 3D-SIM^2^ analysis of HeLa cells immunostained with anti-NuMA (green) and anti-PCNT (magenta) during metaphase and anaphase in the absence and upon acute treatment with MLN8237. Insets on the right show the pole localized NuMA (grey) and PCNT (magenta). (G) Domain organization of AcGFP-tagged and siRNA-resistant wild-type full-length NuMA (**i**) and various deletion/mutant constructs (**ii-v**). (**H, I**) IF analysis of HeLa cells expressing all NuMA constructs mentioned on the left in panel H, after transfections with NuMA siRNA. The DNA is shown in blue. See also figs. S4 and S5. Scale bars in (A), (E), and (H) 10 *μ*m, and in (F) 5 *μ*m for the cell and 2 *μ*m for the insets. Error bars indicate SD.

Next, to monitor the organization of endogenous NuMA protein at the poles in the presence of and upon acute Aurora-A inhibition, we utilized a three-dimensional (3D) lattice-structured illumination super-resolution imaging (3D-SIM^2^). This analysis revealed that NuMA organizes into an extended ‘meshwork’ at the poles during metaphase and anaphase and transformed into highly compact structures upon Aurora A inhibition (Fig. 2F). These experiments suggest that Aurora A activity during mitosis keeps NuMA pools at the poles in dynamic ‘meshwork-like’ assemblies. In the absence of that, NuMA undergoes a phase transition into a non-dynamic compact state, which, from now onwards, we refer to as a ‘solid.’

### The Dynein binding and self-oligomerization capability of NuMA is critical for its pole accumulation

The pole localization of NuMA during metaphase and its altered dynamic state upon Aurora A inhibition recently has been linked to its ability to undergo liquid-liquid phase separation (LLPS) (Sun et al., 2021; Ma et al., 2022). Notably, these conclusions were made: 1) in conditions where GFP-tagged NuMA fragments were overexpressed in the presence of endogenous protein, 2) using recombinant NuMA at high concentration in the presence of crowding reagents, and 3) using aliphatic alcohol 1,6-hexanediol. These experimental conditions can lead to misinterpretation in the context of LLPS, as discussed extensively (McSwiggen et al., 2019; Alberti et al., 2019; Duster et al., 2021; Poudyal et al., 2023; Hedtfeld et al., 2024). Therefore, we first sought to investigate the mechanisms of NuMA accumulation at the poles, followed by addressing Aurora A-guided NuMA phase transition *in vivo*.

Sun et al. found that NuMA tagged with mClover assembled into multiple ‘condensates- like’ bodies that occasionally fuse upon nocodazole treatment (Sun et al., 2021). This observation allowed authors to conclude that NuMA assembles into liquid-like condensates *in vivo* without microtubules and associated dynein/dynactin. In contrast, we found that the concentration of nocodazole (100 ng/ml = ∼332 nM) utilized by the authors is insufficient to depolymerize microtubules fully (Fig. S5 A, B). When microtubules are fully depolymerized using a high concentration of nocodazole (>1.5 mM), NuMA does not assemble into phase-separated condensates *in vivo* (Fig. S5 A, B). These results indicate that NuMA’s ability to accumulate at the poles cannot be explained by its potential to undergo LLPS without microtubule and possibly dynein/dynactin motor interaction.

The accumulation of NuMA at the poles is hypothesized to be regulated by active dynein/dynactin-mediated transport of NuMA (Radulescu and Cleveland, 2010; He et al., 2023). Therefore, to test if NuMA requires the dynein/dynactin motor for its accumulation at the poles *in vivo*, we tested the localization of AcGFP-tagged NuMA lacking the N-terminal (1-705 amino acids) dynein/dynactin interaction module (Kotak et al., 2012) (AcGFP-NuMA^r^ΔDBD). AcGFP- NuMA^r^ΔDBD failed to accumulate significantly at the poles (Fig. 2 G-I; **ii** *vs.* **i;** see Fig. S5 C-E for NuMA depletion). Once at the poles, NuMA should engage in multivalent interactions for its accumulation. Recombinant full-length NuMA can assemble into higher-order multimeric assemblies through its coiled-coil and C-terminal domains *in vitro* (Harborth et al., 1999); thus, we reasoned that synergy between the coiled-coil region and the C-terminus may promote NuMA accumulation at the poles. To this end, we investigated the localization of AcGFP-tagged mutant NuMA lacking most of its coiled-coil domain, except 213-705 amino acid residues, which are required for dynein/dynactin interaction (Kotak et al., 2012; Okumura et al., 2018; Renna et al., 2020) (AcGFP-NuMA^r^NC; Fig. 2G) . Notably, AcGFP-NuMA^r^NC accumulated significantly weaker at the poles (Fig. 2 H, I; **iii** *vs.***i**). Next, we examined the relevance of the C-terminus of NuMA (1700-2115 amino acids) for its accumulation at the poles. NuMA C-terminus is largely disordered (Fig. 3A: identified using https://mobidb.bio.unipd.it/; AlphaFold Protein Structure Database; Necci et al., 2021; Banani et al., 2017), and because of this, we hypothesized that it might engage in homotypic multivalent weak interactions (see the next section). As expected, the expression of AcGFP-NuMA^r^(1-1699) lacking the C-terminal intrinsically disordered region (IDR) failed to accumulate at the poles (Fig. 2 G-I; **iv** *vs.* **i**). Similarly, cells expressing AcGFP- NuMA^r^(1700-2115), containing only C-terminus IDR, had a significantly reduced AcGFP signal at the poles (Fig. 2 G, I; **v** *vs.* **i**). These experiments indicate that the synergy between N-terminal dynein binding, a large coiled-coil domain, and C-terminus IDR is critical for robust NuMA accumulation at the poles.

**Fig. 3.**
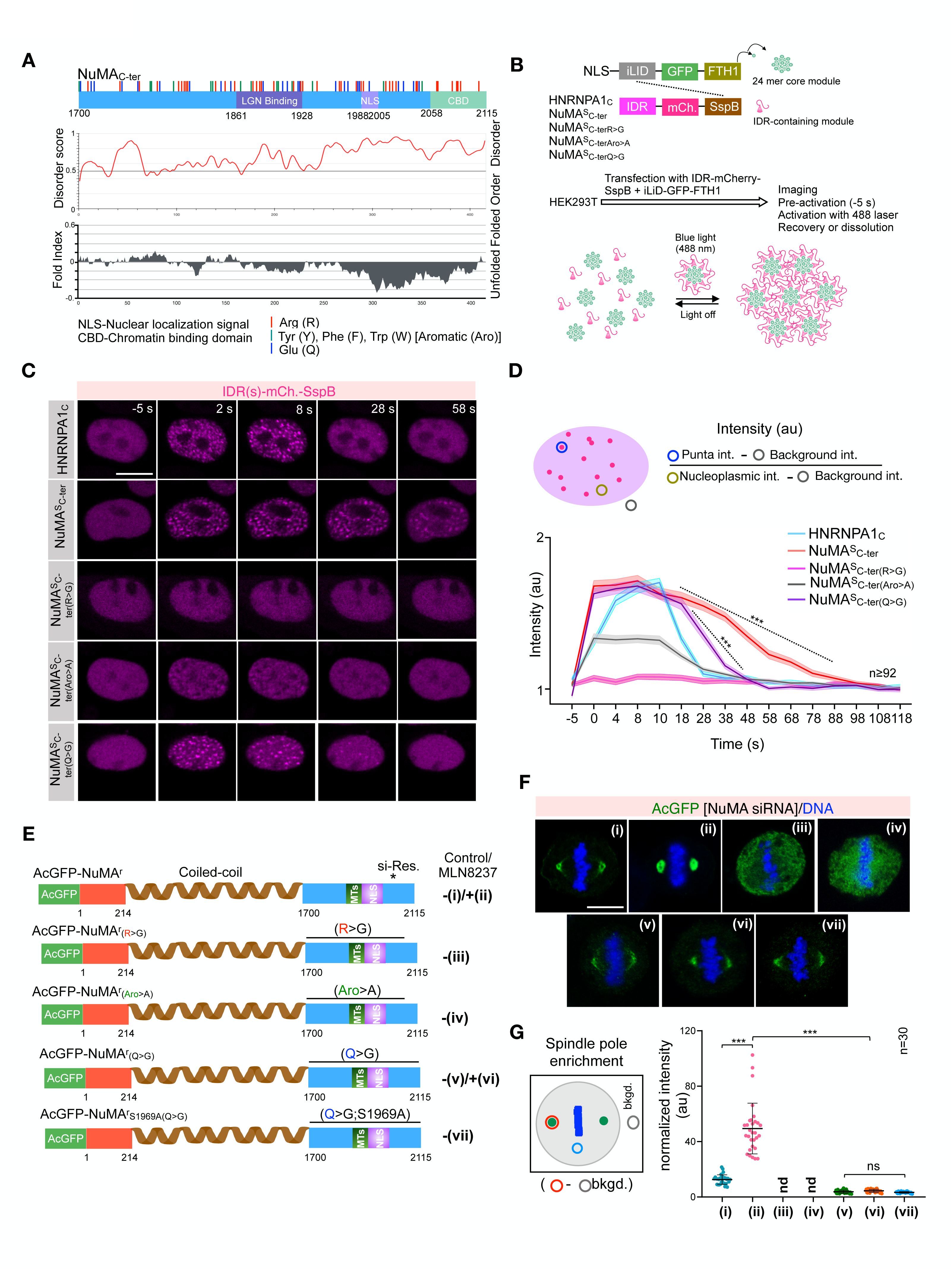
**NuMA’s IDR mediated multivalent interactions are essential for its pole accumulation** (A) Domain organization of NuMAC-ter and the distribution of arginine (R; red), tyrosine, phenylalanine, tryptophan (Aro; green), and glutamine (Q; blue) residues within the NuMAC-ter. The chromatin binding domain (CBD), LGN binding region, and nuclear localization domain (NLS) are marked. IU Pred, intrinsic disorder prediction (https://iupred3.elte.hu/); FOLD, folding prediction using the PLAAC website (http://plaac.wi.mit.edu). (B) Schematic representation of the Corelet system. Corelet consists of two modules: 1) GFP-tagged ferritin core (24 mer), which contains photo-activatable iLID domains, and 2) self-interacting intrinsic disorder region (IDR), mCherry-tag, and light-sensitive iLID binding partner SspB (shown by the dashed line). Upon illumination with a 488 nm laser, up to 24 IDR modules are captured by each Core, which assembles into condensates via multivalent IDR interactions in a reversible light-dependent manner. Note that light- dependent multivalent interactions of NuMAC-ter IDR was studied without chromatin binding domain (CBD; referred to as NuMA^S^C-ter). (C) Panels from time-lapse movies of Corelet-expressing HEK293 cells with various IDRs, as indicated. (D) Schematic representation of the quantification method, intensity (in au), and the dynamics of Corelet- based condensates of HNRNPA1c, NuMA^S^C-ter, and NuMA^S^C-ter mutant fragments. Curves and shaded areas indicate mean ± SEM. (E) Domain organization of AcGFP-tagged and siRNA-resistant wild-type full-length NuMA without (**i**) and with MLN8237 treatment (**ii & vi**), as well as various IDR-based mutant constructs (**iii-vii**), as indicated. (F) IF analysis of HeLa cells expressing all NuMA constructs mentioned in panel E, after transfections with NuMA siRNA. The DNA is shown in blue. (G) Schematic representation of the quantification method and the spindle pole intensity of various NuMA constructs. Note that AcGFP-NuMA^r^(R>G) **(iii)**, and AcGFP-NuMA^r^(Aro>A) **(iv)** do not localize to the poles, and therefore their intensity at the poles is not determined (nd). Error bars indicate SD. Scale bars 10 *μ*m.

### Arginine and aromatic residues govern multivalent homotypic interactions, and glutamine residues promote the dynamic-to-solid phase transition

Since IDR sequences in proteins promote weak interactions (Banani et al., 2017; Holehouse and Kragelund, 2024), we sought to investigate the potential of NuMA’s IDR (Fig. 3A) in engaging multivalent homotypic interactions for its pole accumulation. To investigate this, we used the Corelet system (Bracha et al., 2018). This tool utilizes multivalent 24-mer Ferritin as a ‘core particle,’ which acts as a platform to assemble light-induced phase-separated condensates in the nucleus using IDR-dependent multivalent interactions (Fig. 3B). Because NuMA interacts with DNA with the last 58 amino acids present in its IDR (Rajeevan et al., 2020), and its engagement with the DNA may prevent the light-induced IDR-dependent multivalent interaction, we deleted the last 58 amino acids (referred to as NuMA^S^C-ter), to test IDR-mediated interactions using Corelet system. Importantly, NuMA^S^C-ter formed light-induced condensates analogous to the IDR of HNRNPA1c (Bracha et al., 2018) (Fig. 3 C, D). Next, we sought to investigate the nature of amino acids in IDR facilitating multivalent interactions. The IDR sequence contains 39 arginine (R) and 14 tyrosine (Y), phenylalanine (F), and tryptophan (W) [referred to as Aromatic (Aro)] residues, which are mostly conserved (Fig. 3A; Fig. S5F). These amino acids might engage in cation-π interactions (Wang et al., 2018). To test the role of these residues, we designed two constructs; in one, we mutated all arginine residues in NuMA^S^C-ter, except those in the NLS, to glycine. In the other construct, we mutated all the aromatic residues in NuMA^S^C-ter to alanine. Strikingly, the light-dependent condensate formation of arginine mutated NuMA [NuMA^s^C-ter(R>G)] via the Corelet system is entirely abrogated (Fig. 3 C, D). Similarly, we found that mutation of aromatic residues significantly affected the IDR-mediated oligomerization (Fig. 3 C, D). Next, we examined the relevance of these residues for NuMA accumulation at poles in mitosis. We examined the localization of the full-length AcGFP-tagged NuMA^r^, NuMA^r^(R>G), and NuMA^r^(Aro>A) (Fig. 3E). Interestingly, unlike the wild-type AcGFP-NuMA^r^, AcGFP-NuMA^r^(R>G) and AcGFP- NuMA^r^(Aro>A) failed to localize at the poles (Fig. 3 F, G; **iii** & **iv** *vs.* **i**), suggesting collective interactions between the arginine and aromatic residues in the IDR promote NuMA pole accumulation.

Aurora A phosphorylates NuMA at S1969 in its IDR (Kettenbach et al., 2011). Cells expressing mutant NuMA containing the S1969A replacement strongly accumulate at the poles (fig. S3E), indicating that Aurora A-based phosphorylation at S1969 keeps NuMA in a dynamic state. We reasoned that in the absence of Aurora A-mediated phosphorylation, NuMA IDR undergoes conformational changes, forcing NuMA to adopt a solid configuration. If this assumption is correct, which amino acids would govern NuMA phase transition from dynamic to solid upon Aurora A inhibition? Glutamine (Q) residues promote the hardening of proteins associated with neurodegenerative pathologies (Wang et al., 2018; Patel et al., 2015; Peskett et al., 2018). Thus, we investigated the relevance of glutamine residues in NuMA IDR in facilitating solid configuration upon Aurora A inhibition. Surprisingly, we found no difference in the condensate assemblies of mutant NuMA where glutamine residues are replaced with glycine in the Corelet system; however, the mutant NuMA IDR was relatively more dynamic and dissolved significantly faster than the wild-type IDR (Fig. 3 C, D). To characterize the relevance of glutamine residues for pole localization in mitosis, we examined the localization of either AcGFP-tagged NuMA where all the glutamine in its IDR were mutated to glycine (NuMA^r^(Q>G); Fig. 3E). Notably, we found that AcGFP-NuMA^r^(Q>G) localizes to the poles, albeit significantly less than the wild-type AcGFP-NuMA^r^ (Fig. 3 F, G; **v** *vs.* **i**). Also, the weakly localized AcGFP-NuMA^r^(Q>G) failed to enrich robustly at the poles in cells treated with Aurora A inhibitor MLN8237, compared to AcGFP-NuMA^r^ (Fig. 3 F, G; **vi** *vs.* **ii**). Similarly, the AcGFP-NuMA^r^(Q>G) construct consisting of a replacement of S1969 to non-phosphorylatable alanine [AcGFP-NuMA^r^S1969A(Q>G)] did not significantly accumulate at the poles (Fig. 3 F, G; **vii**). The subtle enrichment of AcGFP- NuMA^r^(Q>G) at poles upon Aurora A inhibition could not be because of its weak localization, as cells that stably expressed low amounts of AcGFP-NuMA^r^ strongly enrich AcGFP-NuMA^r^ at the poles upon Aurora A inhibition (Fig. S5 G, H). Thus, we conclude that glutamine residues in NuMA IDR sequence promote hardening and transform NuMA from a dynamic state into a solid state without Aurora activity.

### Artificially increased multivalent interactions between NuMA molecules mimic Aurora A inactivation

Our data suggest that Aurora A phosphorylation at S1969 on NuMA may prevent strong homotypic multivalent interactions between NuMA molecules. If this hypothesis is true, artificially increasing NuMA multivalency at the poles should lead to a similar outcome as upon Aurora A inhibition. To test this hypothesis, we made a fusion construct between NuMA and Kaede. Kaede is a homotetrameric protein with a size of 116 kDa (Dittrich et al., 2005). We reasoned that a fusion of Kaede with NuMA-which itself is capable of assembling into dimer/multimer using its coiled-coil and IDR would lead to the formation of higher-order multiprotein assemblies of Kaede-NuMA at the poles, somewhat analogous to cells that are inactivated for Aurora A kinase (Fig. 4A). As hypothesized, Kaede-NuMA fusion proteins were significantly enriched at the poles during metaphase and anaphase (Fig. 4B). Kaede-NuMA fusion protein complexes were also detected at the poles during the late anaphase and G1 phase of the cell cycle, in contrast to monomeric AcGFP-tagged NuMA (Fig. 4B). Notably, Kaede-NuMA accumulated at poles had the potential to bend the chromosome ensembles and nascent nuclei around it, similar to those seen upon Aurora A inhibition.

**Fig. 4.**
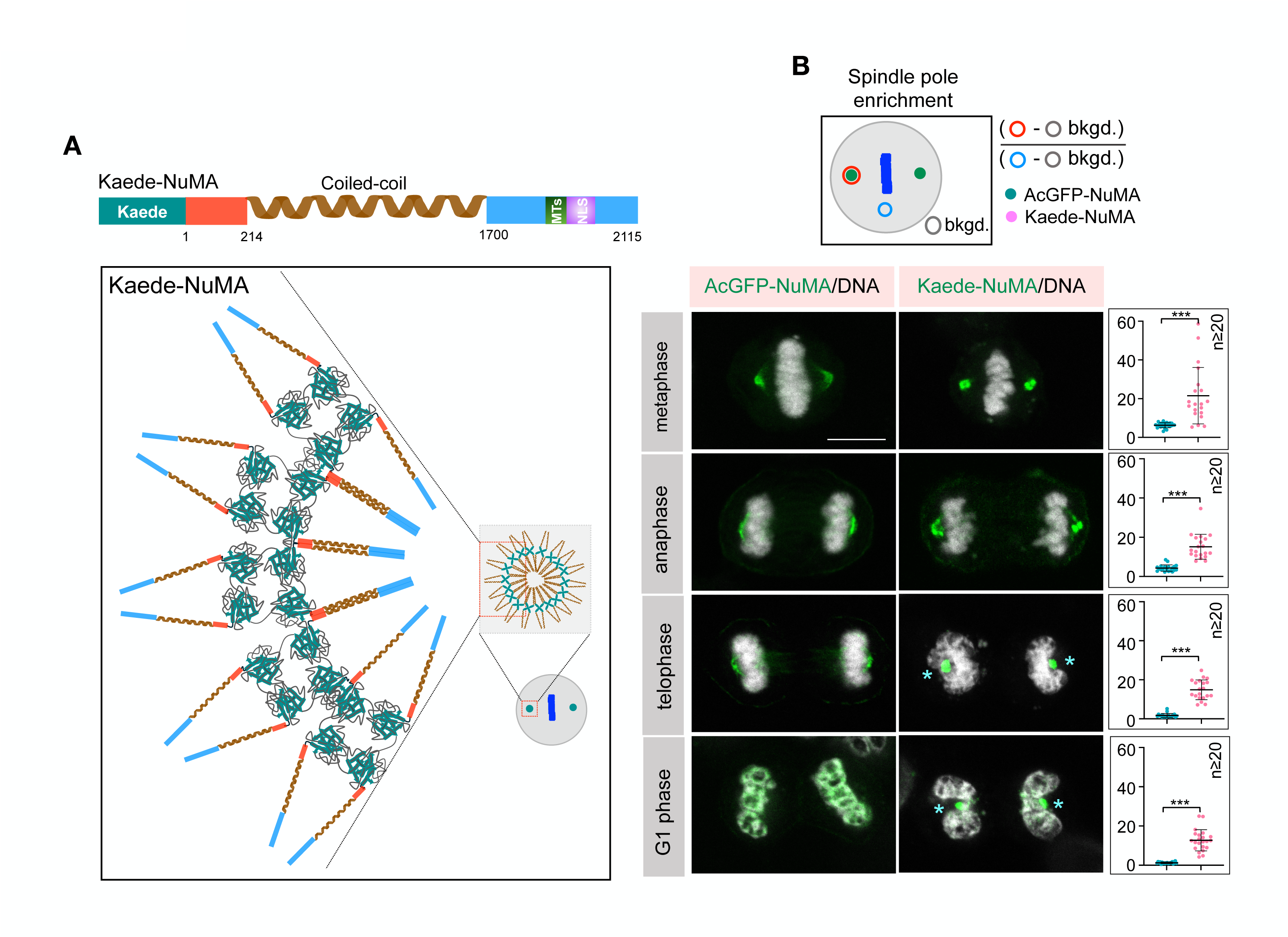
**Forcing multivalent interactions are sufficient to mimic Aurora A inactivation** (A) Schematic depiction of Kaede-tagged full-length NuMA and hypothesized oligomeric assemblies. (B) Schematic representation of the quantification method and the fluorescence intensity measurement at poles in cells expressing AcGFP-NuMA or Kaede-NuMA (in green) during various stages of cell cycle, as indicated. The spindle pole accumulation of AcGFP-NuMA and Kaede-NuMA is quantified on the right for all these stages. DNA is shown in grey. Cyan asterisks depict the bending of chromosome ensemble mass/ nucleus around the Kaede-NuMA poles. Error bars indicate SD. Scale bar 10 *μ*m.

### Proper nuclear morphology at mitotic exit is crucial for nucleoli organization and rDNA decompaction

The impact of Aurora A inhibition on nuclear morphology may influence genome organization at early G1. To evaluate this, we first analyzed the arrangements of nucleoli on the segregated genomes post-nuclear envelope reformation. The chromosomal regions known as nucleolar organizing regions, which are constituted of tandem repeats of ribosomal DNA sequences (rDNA), form the backbone of nucleoli. We reasoned that a potential impact of nuclear morphology on 3D genome organization upon Aurora A inactivation could influence their spatial organization. We followed nucleolus dynamics in cells coexpressing the nucleolus marker-AcGFP-Fibrillarin, mCherry-H2B, and SNAP-NuMA. The AcGFP-Fibrillarin spots were uniformly distributed along the long axis of the newly formed nuclei in control cells (Fig. 5 A, B). In contrast, in MLN8237 treated cells, the nucleoli are considerably more unevenly distributed (referred to as random) (Fig. 5B)

**Fig. 5.**
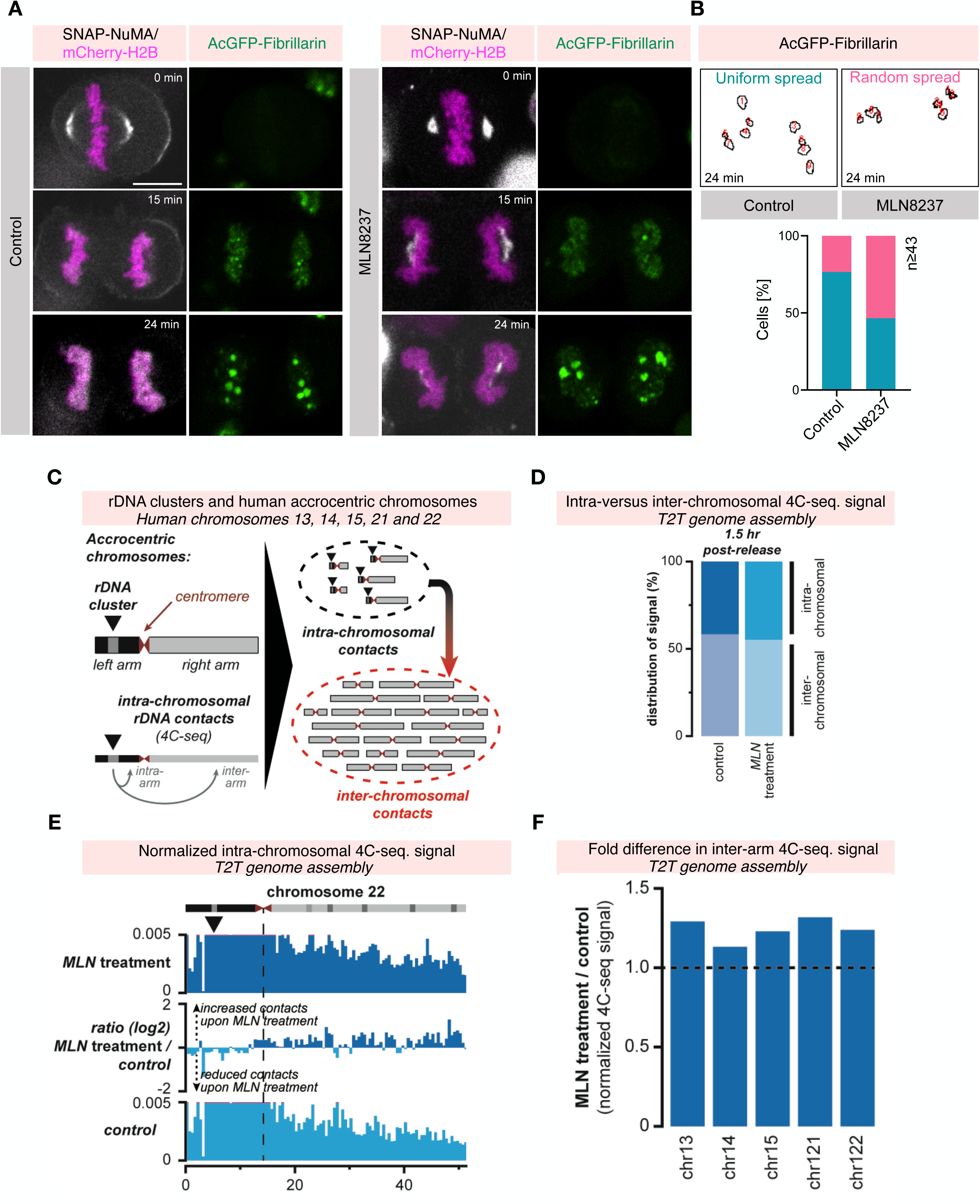
**Abnormal accumulation of NuMA at the poles impacts the organization of nucleoli and rDNA chromosome contacts** (A) Panels from time-lapse movies of HeLa cells coexpressing AcGFP-Fibrillarin (green), mCherry-H2B (magenta) and SNAP-NuMA (grey) in control or treated with MLN8237. Timepoint t=0 min was set to the metaphase-to-anaphase transition. Scale bar 10 *μ*m. (B) Analysis of AcGFP-Fibrillarin foci which were either categorized as uniformly arranged (turquoise), or randomly spread (pink) along the long-axis of the segregated chromosomes. (C) Left: schematic outline of the acrocentric human chromosomes and the location of the rDNA clusters on the short, left chromosomal arms. Below, the analysis of intra-chromosomal contacts using the rDNA clusters as 4C-seq viewpoints. Right: intra-chromosomal versus inter-chromosomal contacts of the rDNA clusters using 4C-seq. (D) Quantification of the combined 4C-seq signal on the five acrocentric chromosomes (intra-chromosomal signal) versus signal on the other chromosomes (inter-chromosomal signal) for cells with and without Aurora A inhibition (MLN8237 treatment) at 1.5 h after mitotic release. (E) Examples of normalized intra-chromosomal 4C-seq signal for the rDNA clusters on the acrocentric chromosomes 13 and 22. Top panels show signal upon MLN8237 treatment and bottom panels signal in control cells. The difference between MLN8237 treatment and controls is indicated in-between (log2 ratio). Black arrows below the chromosome ideograms indicate the centre of the rDNA clusters where the 4C- seq viewpoints are located. (F) Quantification of differential inter-arm contacts upon MLN8237 treatment or control conditions for individual chromosomes. Inter-arm contacts are consistently increased upon MLN8237 treatment after mitotic release, indicative of a delay in chromatin decompaction.

To further scrutinize if redistribution of nucleoli upon acute Aurora A inhibition influenced their genomic surroundings, we performed high-resolution 4C-seq (Circular Chromosome Conformation Capture followed by sequencing; Simonis et al., 2006) using a viewpoint that is common to all rDNA clusters on the five acrocentric human chromosomes (Fig. 5C). Mapping of interactions revealed that MLN8237-treated and control cells engaged to a similar degree in interactions with the non-acrocentric chromosomes that do not contain rDNA clusters (Fig. 5D; Fig. S6 A, B; intra-chromosomal versus inter-chromosomal 4C-seq signal). The uneven distribution of nucleoli along the long axis of the newly formed nucleus is, therefore, not accompanied by a major repositioning of chromosomes relative to each other. Instead, the distribution of intra-chromosomal contacts revealed a noticeable difference. Focusing on intra-arm versus inter-arm contacts, we observed an increased tendency for MLN8237-treated rDNA clusters to contact the right arms on the acrocentric chromosomes (i.e., the arms that do not carry the rDNA tandem repeats) (Fig. 5 D-F; Fig. S6 C-F). Upon inactivation of Aurora A activity, the increased random distribution of nucleoli at 1.5 h after mitotic exit is accompanied by an increase in inter-arm chromosome contacts, suggesting that the chromosomes remain in a more compacted state. These experiments demonstrate that disrupted Aurora A function during mitotic exit and the associated perturbed nuclear morphology lead to altered reestablishment of interphase genome organization.

## Discussion

Defects in nuclear organization during mitotic exit can severely disrupt gene regulation pathways in daughter cells. This study demonstrates that Aurora A activity during anaphase is critical in keeping pole-localized NuMA in a dynamic state (Fig. 6A). In the absence of Aurora A activity, NuMA undergoes a phase transition from a dynamic to a solid state, resulting in abnormal accumulation at the poles (Fig. 6B). These abnormal poles cause the segregated chromosome sets to bend, leading to disorganization of the forming nuclei during mitotic exit (Fig. 6). In human cells, NuMA is present at concentrations below 1 µM (Hein et al., 2015), and its accumulation at the poles depends on motor dynein/dynactin and homotypic multivalent interactions involving its coiled-coil domain and C-terminal IDR. Since IDR-based intermolecular interactions are typically weak (Banani et al., 2017; Holehouse and Kragelund, 2024), our findings suggest that the structured coiled-coil region stabilizes weak cation-π interactions between IDRs. We propose that this may be a common mechanism for stabilizing multi-oligomeric assemblies of many proteins that undergo phase separation via IDR-mediated weak interactions.

**Fig. 6.**
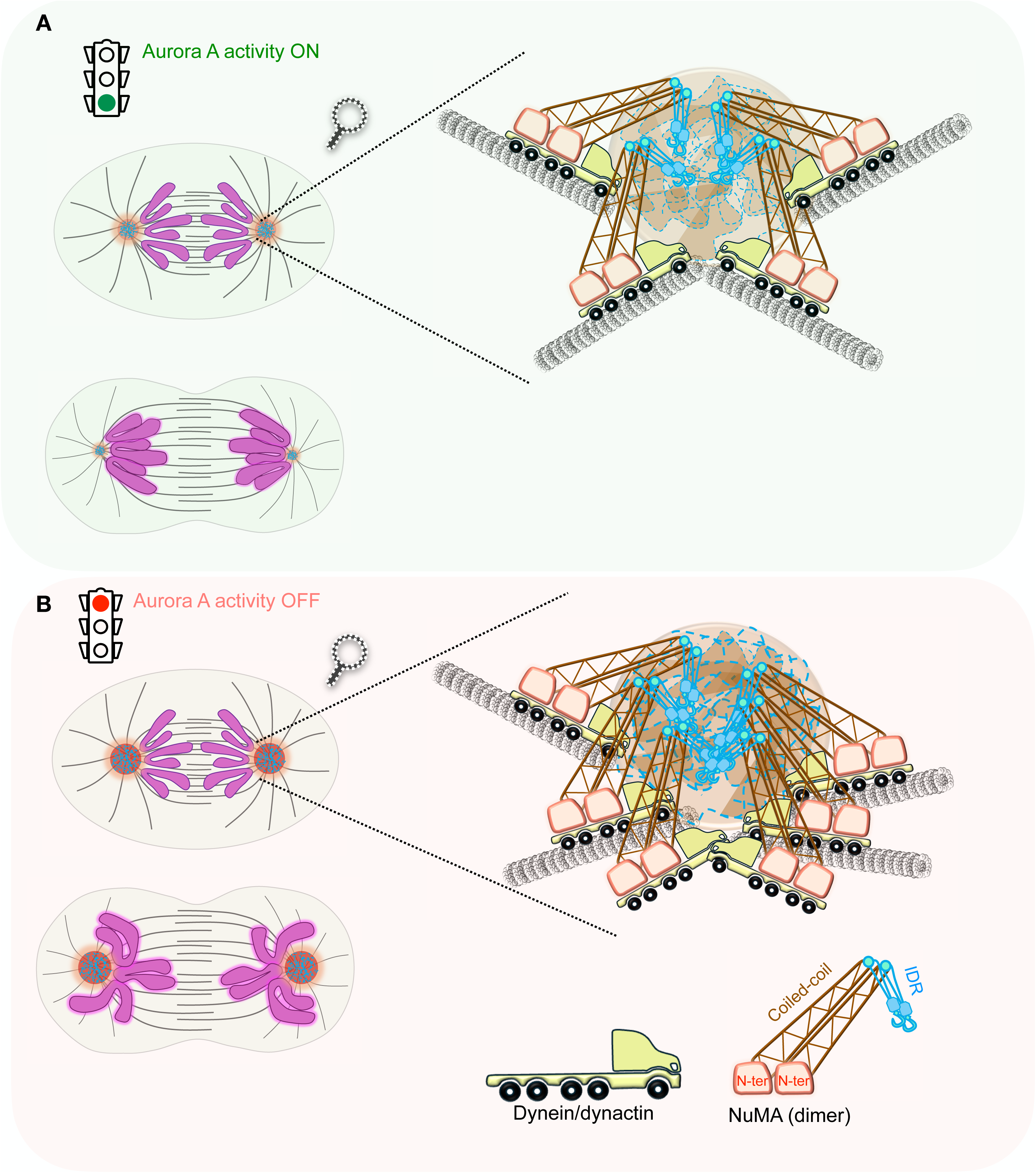
**Aurora A regulates the phase transition properties of NuMA at poles to ensure proper organization of developing nuclei** (A) Model for the coordination between accumulation of NuMA at the poles and chromosome organization at the mitotic exit. In unperturbed control cells, centrosomal Aurora A activity keeps NuMA at the pole in a dynamic state to ensure that NuMA levels at the poles are gradually decreased so that the mitotically segregated chromosome ensemble can form nascent nuclei without any physical hindrance by the pole in late anaphase. Inset highlights that dynein/dynactin (depicted as trucks) carry a dimer of NuMA (shown as cranes) to the poles where the intrinsically disordered regions (IDR) present in the C-terminus of NuMA are engaged in weak multivalent interactions, ensuring its localization and dynamicity. (B) In the absence of centrosomal Aurora A activity, NuMA undergoes a phase transition from dynamic to solid and fails to dissolve at the poles. This leads to the wrapping of chromosome ensemble around the non- dissolved NuMA poles and, therefore, change in nuclear organization post mitosis. Inset highlights that in the absence of Aurora A activity, NuMA (cranes) are engaged in strong multimeric interactions at the poles via their IDR sequences, leading to a change in the phase property of NuMA from dynamic to solid. This work reveals that the coordination between poles (membrane-less organelle) and developing nuclei (membrane organelle) is essential for proper nuclear shape organization.

Notably, Aurora A phosphorylates NuMA at serine 1969 within its IDR. Replacing this serine with a non-phosphorylatable alanine results in abnormal NuMA accumulation at the poles, similar to what is observed when Aurora A is inactive. Since post-translational modifications are known to regulate the material properties of proteins that undergo phase transitions, we hypothesize that phosphorylation at S1969 induces conformational changes in NuMA’s IDR, preventing strong multivalent interactions between NuMA’s IDRs at the poles—interactions that could otherwise be pathogenic. Supporting this hypothesis, we show that artificially promoting NuMA multimerization by fusing it with a tetrameric protein, Kaede, is sufficient to cause NuMA accumulation at the poles, mimicking the effects of Aurora A inactivation.

Super-resolution imaging of NuMA reveals that it assembles into a ‘mesh-like’ scaffold. However, when Aurora A is inhibited, these assemblies become more compact. This behavior is reminiscent of the Drosophila pericentriolar material (PCM) protein Centrosomin (Cnn) and its *C. elegans* homolog SPD-5. During mitosis, Cnn (or SPD-5) is phosphorylated by Plk1 at the centrosome, promoting scaffold assembly around centrioles and facilitating centrosome maturation (Conduit et al., 2014; Rios et al., 2024). Thus, we propose that eukaryotic cells have evolved multiple centrosome/pole-localized proteins with an inherent ability to form scaffolds at the centrosome/poles, ensuring proper microtubule nucleation, spindle assembly, and spindle integrity.

Aurora A regulates several mitotic processes, and since it is often amplified in cancers with poor prognoses, it represents a promising target for cancer therapy. Over the past decade, several highly specific Aurora A inhibitors (e.g., alisertib [MLN8237]) have been developed and tested in clinical trials, but with limited success (O’Connor et al., 2019). Our research highlights a novel post-mitotic role for Aurora A in nuclear organization, suggesting that Aurora A inhibitors may have unforeseen post-mitotic effects if not carefully assessed. Therefore, it is essential to gain a deeper fundamental understanding of Aurora A (and other mitotic kinases) before these inhibitors are used in clinical settings to ensure they are both effective and safe for patients.

## Acknowledgments

We thank A. Desai, J. Pines, M. Lowe, P. Guichard, V. Hamel, K. Subramanian, G. Dey, A. Gorvachev for feedback on the manuscript and discussion. We thank S. Sana, R. Keshri, A. Ghosh, and S. Kapoor for their initial contribution to this project. We acknowledge the help of S. Kapoor with the working model.

## Author contributions

Conceptualization: A.R., V.O., M.B., D.C., S.K.

Methodology: A.R., V.O., M.B., D.C., F.C. D.N. S.K.

Investigation: A.R., V.O., M.B., D.C., F.C., D.N., S.K.

Funding acquisition: D.N., S.K. Supervision: D.N., S.K. Writing – original draft: S.K.

Writing – review & editing: A.R., V.O., M.B., D.C., F. C., D. N., S.K.

## Materials and Methods

### Cell lines and culture

The cell lines used in this study were HeLa Kyoto, RPE1, U2OS and HEK293. All these cell lines were cultured in high glucose Dulbecco’s Modified Eagle Medium (DMEM) supplemented with 10% heat-inactivated fetal bovine serum (FBS) and 100 units/ml of an antibiotic solution containing penicillin and streptomycin at 37°C in a humidified 5% CO2 incubator. All the stable cell lines used in this study were generated in HeLa Kyoto cells (a kind gift from Daniel Gerlich, IMBA, Vienna). HEK293 cells were kindly provided by Dr. Amit Singh (Indian Institute of Science, Bengaluru).

### Generation of stable cell lines

HeLa Kyoto cells stably expressing the specified recombinant constructs were generated by transfecting with 4 µg of plasmid DNA suspended in 500 µl of jetPRIME buffer, incubating it for 5 min followed by the addition of 8 µl of jetPRIME transfection reagent and incubation for another 20 min. The mixture was added to the cells plated at 80% confluency in a 10 cm dish. The medium was replaced after 12 hrs of transfection. After 36 h of transfection, DMEM containing puromycin (400 ng/ml) and/or G418 sulphate (0.4 mg/ml) was added for selection. The clones were isolated, analyzed and confirmed by immunofluorescence and western blotting.

### Transient transfection

For transient plasmid transfection, 2 – 4 µg of plasmid DNA was mixed with 400 µl of serum-free DMEM and incubated for 5 min. After the incubation, 6 µl of Lipofectamine 2000 was added to the mixture and incubated for 15 – 20 min. This mixture was added to the cells plated at 80% confluency on the coverslips or the imaging dish. The cells were fixed or used for immunostaining or live imaging after 24 - 36 h of transfection.

### Small-interfering RNA transfection

For siRNA transfection, 9 µl of 20 µM of the siRNA and 4 µl of Lipofectamine RNAiMAX reagent were mixed with 100 µl of nuclease-free water side-by-side. After 5 min of incubation, these solutions were mixed and incubated for another 20 min. This mixture was added to the cells at 30- 40% confluency on coverslips in 35 mm dish or imaging dish. The medium was replaced after 12 h of siRNA transfection. The cells were grown for a total of 60-72 h followed by fixation in methanol and immunostaining or live-imaging analysis.

### Drug treatments

All the cell lines used in this study were treated with 50 – 250 nM of MLN8237 for Aurora A inhibition for 15 min - 2 h. 500 - 2000 nM nocodazole treatment was given for 17 h for microtubule depolymerization experiments. Following the drug treatments, the cells were analyzed using immunofluorescence, live cell imaging, or western blot analysis.

### Indirect immunofluorescence and live imaging

The cells were grown on autoclaved coverslips (Cat# 40990103, 40990109, Assistant, Germany), and after the transfection with siRNA/ plasmids or drug treatments, they were fixed by cold methanol (-20°C) and incubated at -20°C for 8-10 min. They were washed with PBS and permeabilized with PBST (PBS with 0.05% Triton-X 100) for 5 min, followed by blocking with 1% BSA (in PBST) for 1 hr. Then, the cells were incubated with appropriate primary antibodies, diluted in 1% BSA for 4 h and washed thrice with PBST at 5 min intervals. Following this, the cells are incubated with fluorophore-labeled secondary antibody for 1 h and washed thrice with PBST at 5 min intervals. Further, the cells were incubated with 1µg/µL Hoechst 33342 for 5 min, followed by three PBST washes. After that, the coverslips were mounted using Fluoromount-G. The images were acquired by an Olympus FV 3000 confocal laser scanning microscope using a 60X (NA 1.4) oil immersion objective. The acquired images were analyzed and processed using ImageJ software, preserving the relative intensities.

The time-lapse live imaging of cells plated on an imaging dish (Cat# 0030740017, Eppendorf, Germany) was performed on an Olympus FV 3000 confocal laser scanning microscope using a 40X (NA 1.3) oil immersion objective (Olympus Corporation). The images were captured by the inbuilt FV3000 software at 1, 2, 3 and 5 min intervals, with 9-11 optical sections (3 μm apart). During the imaging, the cells expressing AcGFP/mCherry/SNAP/mEOS tags were maintained at optimal growth conditions (37°C, 5% CO2 and 90% relative humidity) by a Tokai Hit STR Stage Top incubator.

The super-resolution imaging of NuMA and PCNT was done using a 3D-lattice structured illumination super-resolution microscope (3D-SIM^2^) from Zeiss Elyra 7 with Lattice SIM^2^ model with a 60X oil immersion objective.

### Preparation of cell extracts and immunoblotting analysis

The cells were synchronized at prometaphase using 100 nM nocodazole for 17-20 h. For anaphase synchronization, 10 µM of RO-3306 was added to the nocodazole-treated cells for 15-20 min. MLN8237 is added if required. The cells were collected there and suspended in a lysis buffer containing Tris pH 7.4 (50mM), NP-40 (1% v/v), sodium deoxycholate (0.25%), NaCl (150mM), SDS (0.01%), PMSF (0.1 mM) and protease inhibitor cocktail (complete, EDTA-free, 1:1000). The cells were incubated on ice for 1.5 hrs and centrifuged at 13000 rpm for 10 min at 4°C, the supernatant was collected, and protein concentration was quantified by Bradford assay.

The protein samples were normalized to 2 µg/µL using 2X SDS loading dye containing β-mercaptoethanol (4.9%), and the samples were heated at 98°C for 10 min. 20-50 µg of the protein samples were loaded to 6-10% SDS PAGE gels (according to the size of the protein of interest) and resolved in SDS PAGE running buffer (25 mM Tris (pH=8), 192 mM glycine and 0.1% SDS). After electrophoresis, the proteins were transferred to a nitrocellulose membrane using a wet-transfer apparatus at 250 mA for 1.5 h with cold transfer buffer (25 mM Tris (pH=8), 192 mM Glycine). Then, the nitrocellulose membrane was blocked on 5% skimmed milk in 1X PBS-T (0.05% Tween-20) for 1 h, followed by one PBS-T wash. The membrane was incubated in primary antibody (in 5% BSA or 1% skimmed milk) overnight at 4°C. The blot was washed thrice with PBS-T at 5 min intervals, and then a secondary antibody (in 1% skimmed milk) conjugated with HRP was added and incubated at room temperature for 1 hr. Further, the blots were washed thrice with PBS-T at 5 min intervals, and the blots were developed by BioRAD substrate.

### Fluorescence Recovery after Photobleaching (FRAP)

FRAP was performed in HeLa cells stably expressing AcGFP-NuMA and treated with DMSO, MLN8237 (30 min). For the experiment, a pre-bleach image was acquired, followed by a brief pulse of 488 nm laser (40% laser intensity) to bleach the GFP signal from one of the two spindle poles at the metaphase or anaphase stage of the cell cycle, at an approximate area of 10 μm^2^. Post-bleach images were taken every 5 sec for a duration of 50 cycles. All the images were acquired over three z planes (step size = 2 μm, thickness = 6 μm) and the recovery was assessed from the maximum intensity projected images. To account for the fluorescence loss due to imaging-induced photobleaching, fluorescence from the spindle pole separated from the bleached region was simultaneously recorded (control spindle pole). The intensity value in the bleached area was measured and corrected for the background. For metaphase spindle pole FRAP, the curves were normalized using the following equation (Phair and Misteli, 2000):

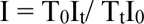

where T0 is the pre-bleach intensity of the control spindle pole, It is the intensity of the bleached spindle pole at timepoint t, Tt is the intensity of the control spindle pole at timepoint t, and I0 is the pre-bleach intensity of the bleached spindle pole.

### Photoconversion experiment

Photoconversion experiments were conducted in the metaphase stage of HeLa cells transiently transfected with mEos-NuMA siRNA background. A circular ROI (∼10 µm^2^) was drawn around a spindle pole and illuminated with a 405 nm laser set at a power of 5% for ∼ 5 sec. A green-to-red photoconversion of the labelled NuMA at the spindle pole was observed, and the exchange was recorded using z-stacking with 3 µm with a 10 sec time interval. The z-stacked images were processed using ImageJ and are represented as maximum z-projections.

### Quantification and statistical analysis

Fiji/ImageJ (https://fiji.sc), GraphPad Prism and Imaris (http://www.bitplane.com/imaris/imaris) were used for all the quantifications. For intensity measurements, the “integrated density” measurement in arbitrary units (au) was used, which is defined as “(sum of pixel values in selection) x (area of one pixel)”. All the quantifications in a given experiment were measured by a constant ROI. The intensity values are plotted by normalizing with the cytoplasmic intensities and corrected for the background as mentioned in the figure.

The level of significance between two mean values was calculated by a two-tailed unpaired Student’s t-test. If the P value is less than 0.05, it is considered to be statistically significant. It was calculated and confirmed using GraphPad prism 8 (http://www.graphpad.com/scientific-software/prism/). The levels of significance are mentioned as n.s. (not significant) if p>0.05, *p < 0.05, **p < 0.01, and ***p < 0.001.

ImageJ was used to measure the line scan intensity using the straight line tool with a width of 2 um. Circularity, solidity, and aspect ratio were also measured in ImageJ.

### Assigning time ‘0’

In almost all experiments, time, t = 0 is assigned as metaphase-to-anaphase transition. This is the frame at which a proper metaphase plate is formed, leading to anaphase entry in the upcoming time frame.

### Optogenetic activation of the Corelet system

HEK293 cells were transiently transfected with 2 µg each of iLid-GFP-FTH1 and IDR(s)- mCherry-SspB at 70% confluency plated on coverslips in 35 mm dish. Cells were imaged 24 h after plasmid transfection using the mCherry (561 nm) channel for the visualization of the behaviour of IDR components. For global or local activation, the cells were illuminated with a 3- 5% blue laser (480 nm) for 1-20 sec in a circular area of 3000 – 4000 µm^2^ and imaged for 2 min with 2-sec intervals with a single z-plane. The puncta intensities were measured using ImageJ (ROI of area 0.893 µm^2^) by taking the ratios of fluorescent intensities at the puncta with respect to the cytoplasm, corrected for the background, and plotted.

### Sample preparation for 4C-seq experiments

For synchronization of HeLa Kyoto cells in the early G1 phase for 4C analysis, 24 h after plating the cells, they were synchronized in prometaphase with 100 nM nocodazole for 17 h. The synchronized cells were collected by mitotic shake-off and washed thrice with 1X PBS to remove nocodazole. Cells were plated in media containing DMSO or MLN8237 for 1.5 h followed by harvesting for 4C analysis. Briefly, 10-15 x 10^6^ cells were collected, washed once with 1X PBS, and cross-linked using freshly prepared 2% formaldehyde solution by tumbling for 10 min at room temperature. The fixation was stopped by adding glycine and immediately transferring the tubes to ice. The cross-linked cells were collected at 300g for 5 min at 4℃ and washed once with 1X PBS. After removing the supernatant, the pellet was snap-freeze in liquid nitrogen and stored at - 80°C.

### 4C-seq experiments and analysis

Chromatin fixation, cell lysis and 4C-seq procedure were done as previously described using 10- 15 x 10^6^ cells per cell experiment (Matelot and Noordermeer, 2016). *MboI* (New England Biolabs) was used as the primary restriction enzyme and *NlaIII* (New England Biolabs) as the secondary restriction enzyme. 4C-seq sequencing libraries using the rDNA viewpoint, which is present in the human genome in several hundred copies, were generated using a 2-step amplification approach with reduced amounts of input, as described in (Haarhuis et al., 2017; Krijger et al., 2020). In the first step, a total of 120ng of 4C material was amplified using the Expand Long Template PCR System (Roche Diagnostics) in 12 reactions in parallel for 12 cycles. All reactions were pooled, followed by clean-up of 1/6^th^ of the material using Agencourt Ampure XP Beads (Beckman Coulter) to remove fragments smaller than 200 bp. In the second step, 20% of the purified PCR products were further amplified in 2 reactions using the Expand Long Template PCR System and universal index adapters for 20 cycles. Amplified 4C-seq libraries were again purified with Agencourt Ampure XP Beads to remove fragments smaller than 200 bp. Quality and size distribution of the PCR libraries was verified by Qubit ds DNA BR kit (Thermo Fisher Scientific) and Tapestation Genomic DNA reagents (Agilent). Amplified 4C-seq libraries from the cells with or without MLN treatment were mixed in an equimolar amount, followed by sequencing using 86bp single-end reads on the Illumina Next-seq 550 device at the I2BC High-throughput sequencing facility (Gif-sur-Yvette, France).

4C-seq datasets were processed using the c4ctus pipeline (Miranda et al., 2022), available at https://github.com/NoordermeerLab/c4ctus. Mapping was done on the T2T-CHM13 genome assembly. For visualization, 4C-seq tracks were binned to 500kb resolution and then each bin was normalized to the total signal for each chromosome individually.

### Sequence analysis for protein disorder

IUPred program (https://iupred3.elte.hu/) was used to analyze the disordered regions NuMA.

### Cloning of different constructs

Different truncated constructs of NuMA (AcGFP-NuMA^r^(1-1699), AcGFP-NuMA^r^ΔDBD, AcGFP- NuMA^r^(1700-2115)) were generated by amplifying the regions (Fig. 2H) from AcGFP-NuMA^r^ by respective primer pairs and subcloning them with AgeI and NotI restriction sites inside pIRES-AcGFP-FLAG vector.

AcGFP-NuMA^r^ NC was generated by sequential cloning where initially, amino acids 1-705 of NuMA were subcloned using AgeI and EcoRI sites. After that, amino acids 1700-2115 were subcloned using EcoRI and NotI sites inside the pIRES-AcGFP-FLAG vector. As a result of sequential cloning, the EF linker was incorporated at the junction of these two fragments.

AcGFP-NuMA^r^ (WT),(R>G),(Q>G),(Aro>A) constructs were also made by sequential cloning where in the first round 1-1699 region of NuMA was subcloned using AgeI and EcoRI sites and in the next round 1700-2115 regions of WT and mutated NuMA (R>G, Q>G, Aro>A) were subcloned using EcoRI and BamHI sites inside pIRES-AcGFP-FLAG vector. The mutated 1700-2115 fragments were synthesized by Twist Bioscience (https://www.twistbioscience.com/) and were subcloned by amplifying with proper primer pairs.

While generating the optogenetic constructs of NuMAC-ter, a common vector system was essential, so we made an in-house vector by subcloning mCherry-SspB using AgeI and NotI sites inside H2B-mCherry vector. In this vector, H2B was substituted with NuMAC-ter using XhoI and AgeI sites. Other mutated constructs were made using a similar strategy. HNRNPA1c-mCherry-SspB construct was generated by amplifying HNRNPA1c-mCherry-SspB from addgene plasmid #122668 and subcloned it in addgene plasmid #21044 using NheI and NotI sites.

---

**Fig. S1.**
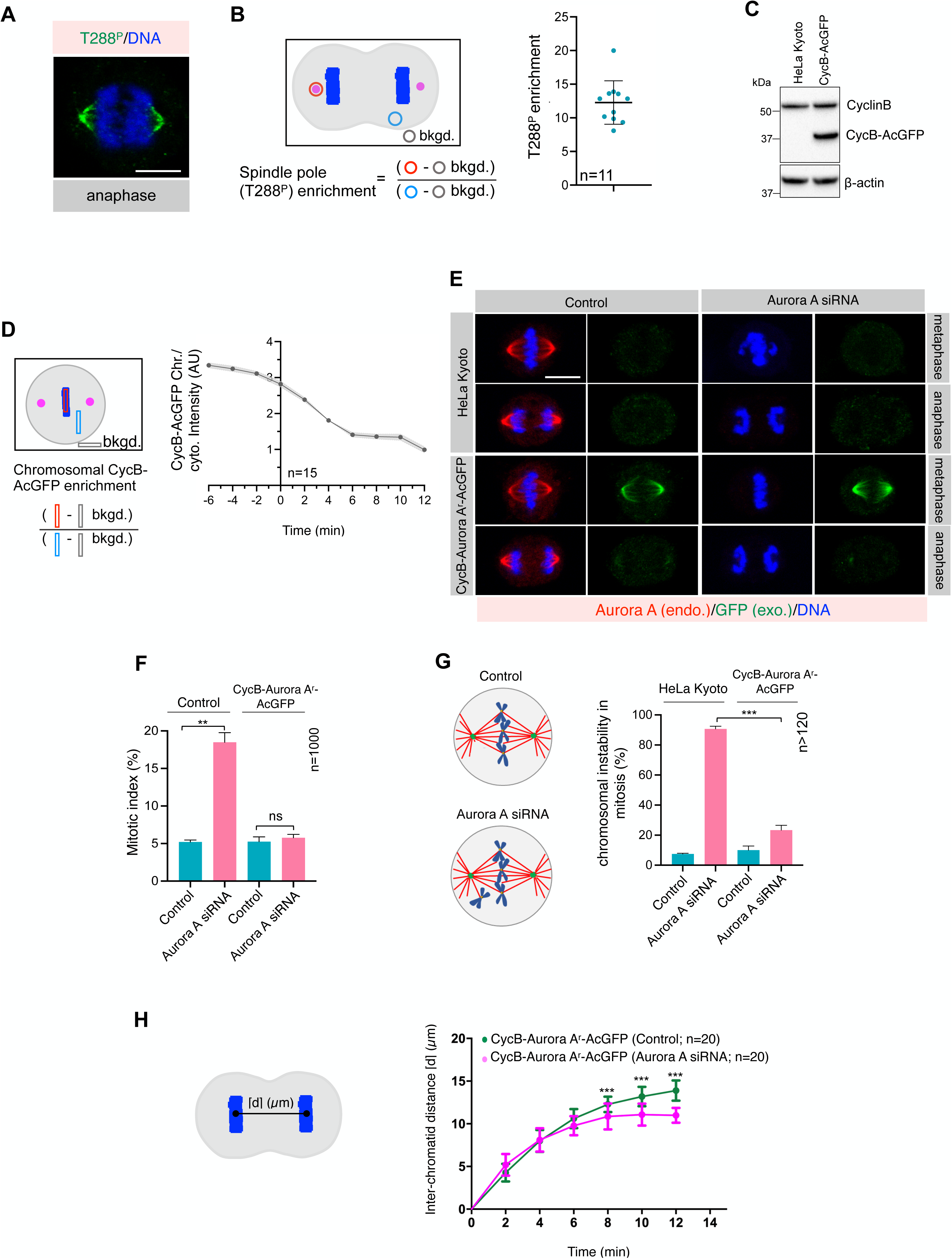
**Anaphase-specific Aurora A degradation tool to study Aurora A function during mitotic exit** (A) Immunofluorescence (IF) analysis of control HeLa cells in anaphase that are stained with phospho-specific anti-Aurora A antibody against T288 (T288^P^; green). DNA is shown in blue. (B) Schematic representation of the quantification method and the quantification of T288^P^ at poles in cells stained with anti-T288^P^ antibody in anaphase. Note that T288^P^ is significantly enriched at the poles in anaphase, suggesting Aurora A is active during anaphase. Error bars indicate SD. (C) Immunoblot analysis of protein extracts prepared from mitotically synchronized control HeLa or HeLa cells stably expressing CycB-AcGFP. Extracts were probed with antibodies directed against CyclinB and b- actin. Endogenous CyclinB and CycB-AcGFP bands are indicated. In this and other immunoblot panels, the molecular mass is indicated in kilodaltons (kDa) on the left. (D) Schematic representation of the method and the quantification of the chromosomal intensity of CycB- AcGFP (in au) during metaphase-to-anaphase transition (time 0). bkgd., representing background intensity. (E) IF analysis of control and monoclonal cell line expressing CycB-Aurora A^r^-AcGFP. These cells are stained with anti-Aurora A (red; for endogenous protein detection) and anti-GFP (green; for exogenous protein detection) antibodies upon transfection with control and Aurora A siRNA for 60 h. DNA is shown in blue. (F) Quantification of the fraction of mitotic cells (mitotic index) in control cells and monoclonal cell line expressing CycB-Aurora A^r^-AcGFP. Cells were analyzed by IF 60 h after transfection with control and Aurora A siRNA. Bars indicate mean ± SD. In this and other Figures, ns- p>0.05; *-p<0.05; **-p<0.01; ***- p<0.001 as determined by two-tailed Student’s t-test. (G) Schematic representation of chromosome instability defects and the quantification of such defects in control cells and monoclonal cell line expressing CycB-Aurora A^r^-AcGFP upon transfection with control and Aurora A siRNA for 60 h, as indicated. Error bars indicate SD. (H) Schematic representation and the estimation of chromosome separation kinetics of monoclonal cell line expressing CycB-Aurora A^r^-AcGFP in control or upon transfection with control and Aurora A siRNA for 60 h. Error bars indicate SD. Scale bars 10 *μ*m.

**Fig. S2.**
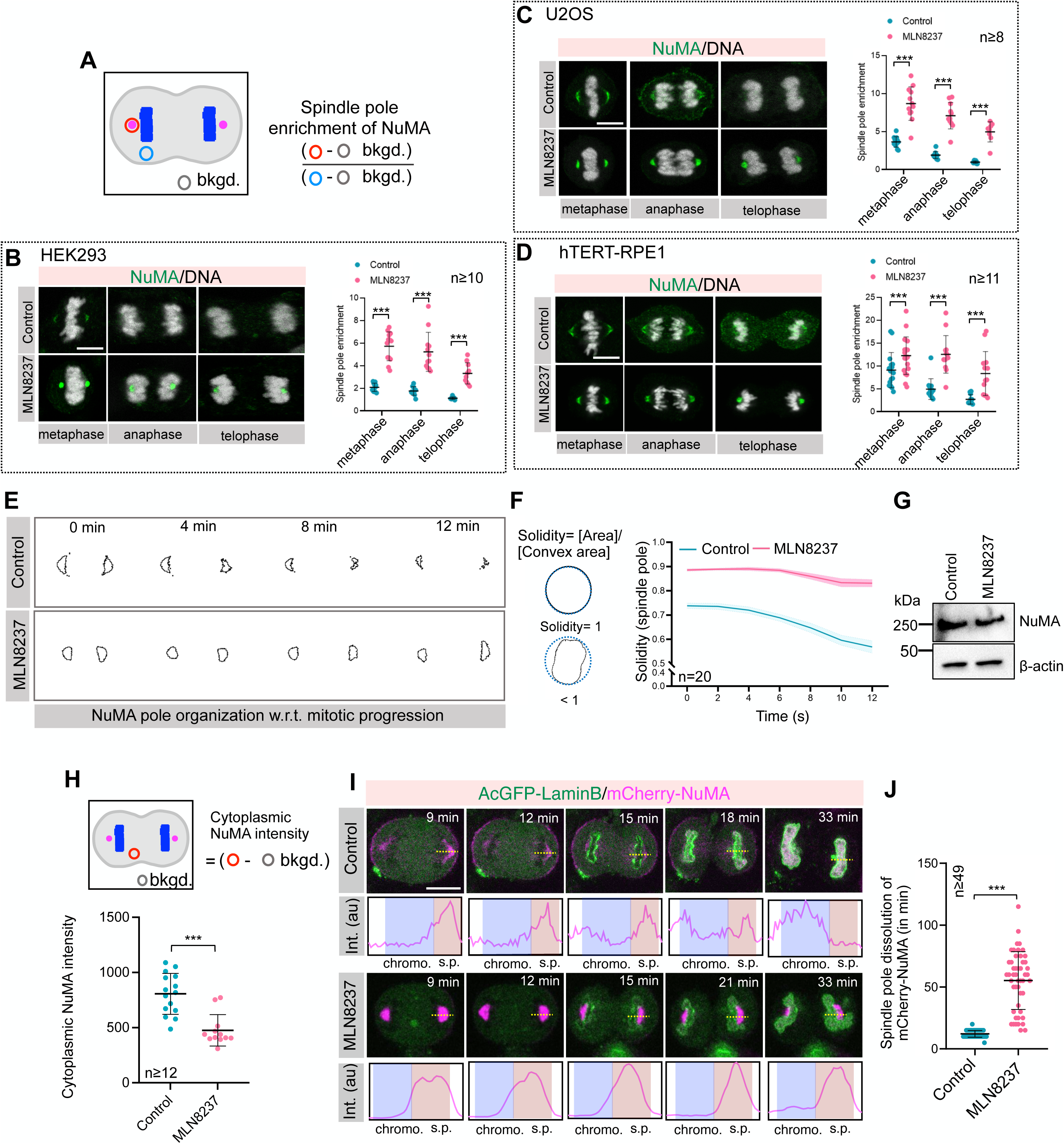
**Endogenous NuMA is significantly enriched at the poles upon acute Aurora A inhibition** (**A-D**) Schematic representation of the method for quantifying spindle pole intensity of NuMA (in au) (A) and the outcome of such analysis by performing IF in various cell lines such as HEK293 (B), U2OS (C), and hTERT-RPE1 (D) in control and upon acute Aurora A inhibition with MLN8237. For this analysis, cells are stained with anti-NuMA (green) antibody. DNA is shown in grey. The quantification represents SD. (E) The representative geometry of NuMA organization at the pole in control and MLN8237-treated HeLa cells. Timepoint t=0 min was set to the metaphase-to-anaphase transition (F) The measurement of the solidity of NuMA-based poles in control and MLN8237-treated cells. Curves, and shaded areas indicate mean ± SEM. The p-values for the solidity of NuMA-based poles for all the time points is <0.001. (G) Immunoblot analysis of protein extracts prepared from anaphase synchronized control and MLN8237- treated cells. Extracts were probed with antibodies directed against anti-NuMA and anti-β-actin. (H) Schematic representation of the method for quantifying cytoplasmic NuMA intensity during anaphase and the outcome of such analysis in arbitrary unit (au). Error bars indicate SD. (I) Panels from time-lapse movies of representative HeLa cells stably coexpressing AcGFP-LaminB1 (green) and mCherry-NuMA (magenta) in the absence (control) or presence of MLN8237. Timepoint t=0 min was set to the metaphase-to-anaphase transition (not shown). Line-scan analysis shows chromosome (chromo.) and poles (s.p.) localized NuMA in a single confocal section. (J) Quantification of the dissolution time (in min) of mCherry-NuMA at the poles in control and MLN8237- treated cells w.r.t. metaphase-to-anaphase transition. Error bars indicate SD. Scale bars 10 *μ*m.

**Fig. S3.**
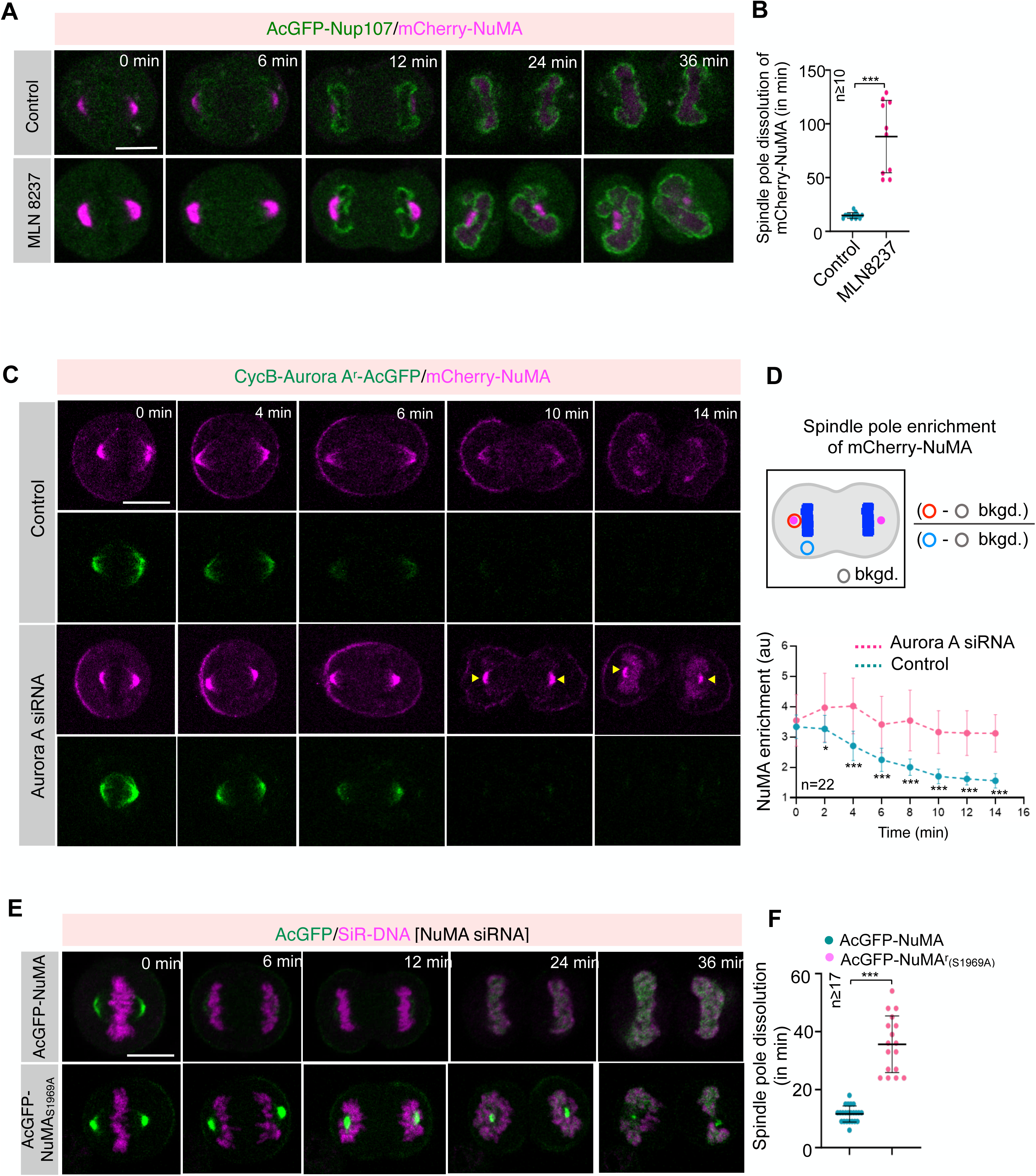
**Organization of the nascent nuclei is markedly affected in cells with abnormal NuMA accumulation at the poles** (A) Panels from time-lapse movies of representative HeLa cells expressing nucleoporin marker AcGFP- Nup107 (green) and mCherry-NuMA (magenta) in the absence (control) or upon acute inhibition of Aurora A kinase using MLN8237. Timepoint t=0 min was set to the metaphase-to-anaphase transition. (B) Quantification of the dissolution time (in min) of mCherry-NuMA at the poles in control and MLN8237- treated cells w.r.t. metaphase-to-anaphase transition (B). Error bars indicate SD. (C) Panels from time-lapse movies of representative HeLa cells stably coexpressing CycB-AuroraA^r^-AcGFP (green) and mCherry-NuMA (magenta) during anaphase in control and upon transfection with Aurora A siRNA. Recording was started 60 h post-transfection for control siRNA and Aurora A siRNA. (D) Schematic representation of the quantification method to analyze spindle pole enrichment of mCherry-NuMA intensity (in au) during anaphase in cells stably coexpressing CycB-AuroraA^r^-AcGFP and mCherry-NuMA in control and after transfection with Aurora A siRNA. bkgd., background intensity. Error bars indicate SD. (E) Panels from time-lapse movies of representative HeLa cells stably coexpressing either AcGFP-NuMA or AcGFP-NuMAS1969A (green) and probed for SiR-DNA (magenta) to visualize chromosomes ensemble during anaphase. Recording was started 60 h post-transfection for control siRNA and NuMA siRNA. (F) The graph represents the quantification of AcGFP-NuMA or AcGFP-NuMAS1969A dissolution time (in min) at the poles w.r.t metaphase-to-anaphase transition (D). Error bars indicate SD. Scale bars 10 *μ*m.

**Fig. S4.**
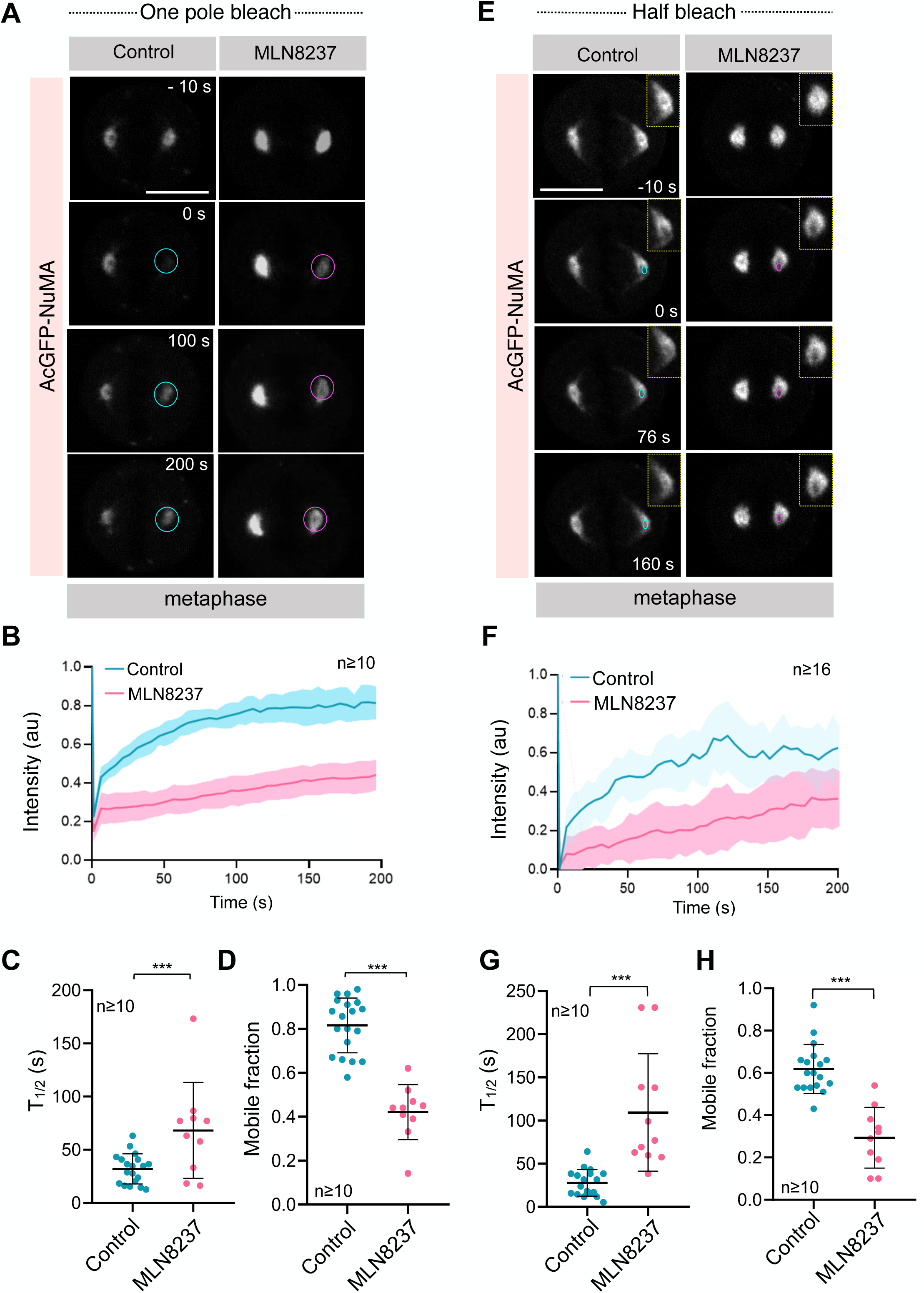
**NuMA undergoes phase transition from dynamic-to-solid state upon Aurora A inhibition** (A) FRAP analysis of metaphase cells stably expressing AcGFP-NuMA (grey) in the absence or presence of MLN8237. Time is indicated in seconds (s). Blue and magenta circles show the bleached regions of control and MLN8237-treated cells, respectively. (B) The AcGFP recovery profile of the bleached area corrected for photobleaching is plotted for 200 s for control and MLN8237-treated cells. Curves and shaded areas indicate mean ± SD. Note the remarkably slow recovery of pole-localized AcGFP-NuMA signal in MLN8237-treated cells. (**C, D**) The half-time of recovery [T1/2] (C) and the mobile fraction (D) of untreated and MLN8237-treated metaphase cells. Error bars indicate SD. (E) Half-FRAP analysis of metaphase cells stably expressing AcGFP-NuMA (grey) in the absence or presence of MLN8237. Time is in (s). Blue and magenta circles show the bleached regions within the photo-manipulated areas of control and MLN8237-treated cells, respectively. (F) The AcGFP recovery profile of the bleached area corrected for photobleaching is plotted for 200 s for control and MLN8237-treated cells. Curves and shaded areas indicate mean ± SD. Note the remarkably slow recovery of AcGFP-NuMA within the photo-manipulated area in MLN8237-treated cells, compared to control cells. (**G, H**) The half-time of recovery [T1/2] (G) and the mobile fraction (H) of untreated and MLN8237-treated Half-FRAP metaphase cells. Error bars indicate SD. Scale bars 10 *μ*m.

**Fig. S5.**
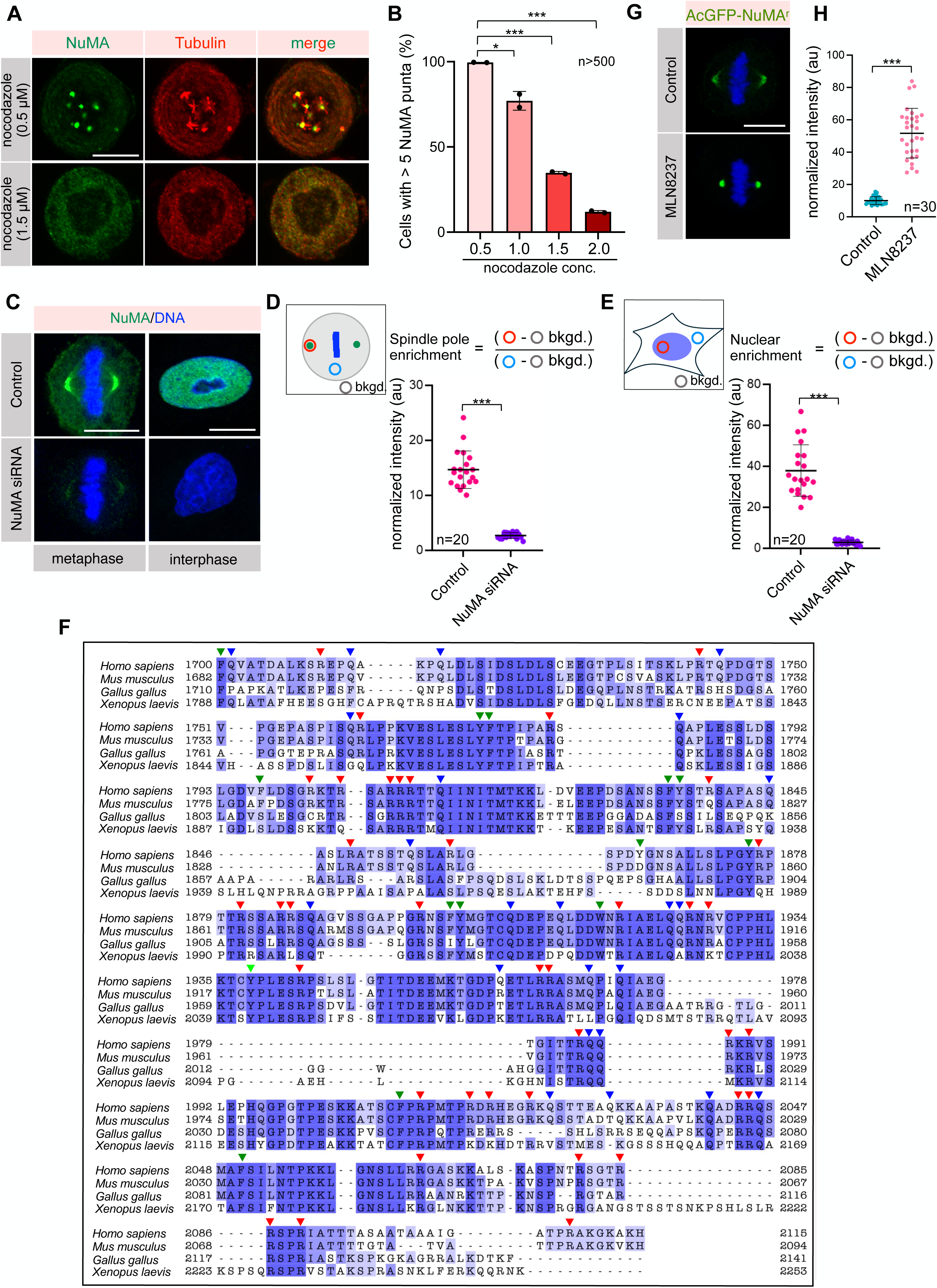
**Microtubules are essential for assembly of NuMA into condensates-like bodies** (**A, B**) IF analysis of HeLa cells after treatment with various concentrations of microtubule poison nocodazole for 17 h. These cells were stained with anti-NuMA (green) and anti-β-tubulin (red) antibodies. The quantification in B depicts the presence of more than 5 NuMA-based condensates at various nocodazole concentration. Note, that NuMA fails to assemble into condensates at high nocodazole concentration (>1.5 *μ*M), suggesting NuMA required microtubules for its accumulation in the condensates-like bodies. Error bars indicate SD. (C) IF analysis to assess the efficiency of NuMA depletion after 72 h of transfection with NuMA siRNA. IF analysis show NuMA localization during mitosis and in interphase. (**D, E**) NuMA quantification at the poles (D) and in the nucleus (E). NuMA was stained using anti-NuMA antibodies (green), and DNA is shown in blue. Error bars indicate SD. (F) Sequence alignment of the NuMA C-terminus (1700-2115 aa) of different NuMA orthologs. Red, green, and blue arrowheads represent arginine (R), aromatic (Aro: W, Y, F), and glutamine (Q) residues, respectively. (G) A representative HeLa cell stably expressing low levels of siRNA resistant AcGFP-tagged NuMA (in green) that are depleted for endogenous NuMA by siRNA. These cells are either left untreated, or treated with MLN8237, as indicated. DNA is shown in blue. (H) Spindle pole intensity of AcGFP-NuMA^r^ at poles in control and MLN8237-treated cells. Error bars indicate SD. Scale bars 10 *μ*m.

**Fig. S6.**
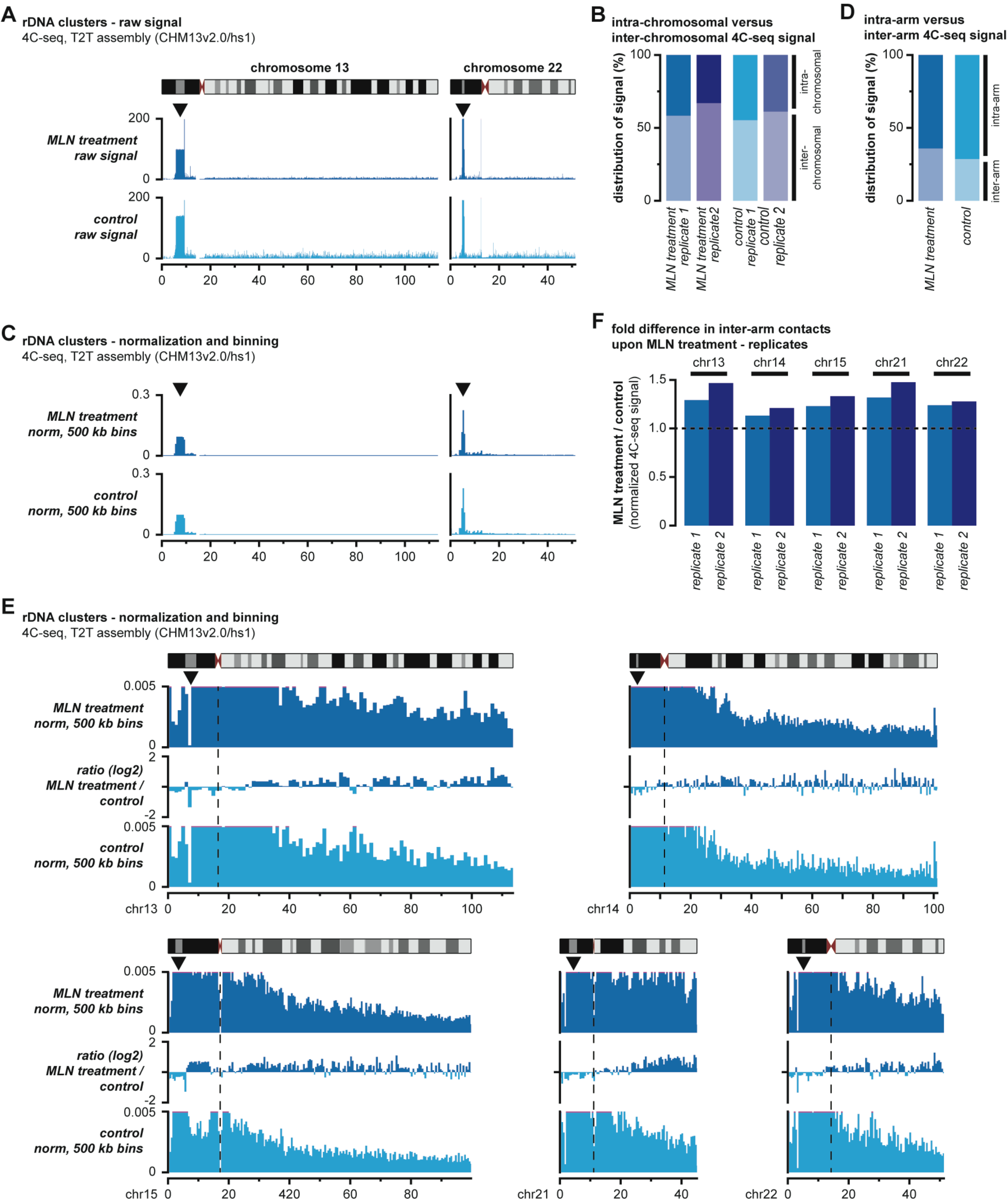
**Reorganization of rDNA contacts upon Aurora A inhibition** (A) Examples of raw intra-chromosomal 4C-seq signal for the rDNA clusters on the acrocentric chromosomes 13 and 22 in MLN8237 treated cells and controls at 1.5 h after mitotic release. Chromosome ideograms are depicted above, with the burgundy arrowheads indicating the position of the centromere and the black arrowheads indicating the centre of the rDNA clusters on the short arm where the 4C-seq viewpoints are located. (B) Quantification of the combined 4C-seq signal on the five acrocentric chromosomes (”intra-chromosomal”) versus the other chromosomes (”inter-chromosomal”) for replicate samples at 1.5 h after mitotic release. (C) Examples of normalized and binned 4C-seq signal at 500 kb resolution for the rDNA clusters on chromosomes 13 and 22. Normalization was performed for each chromosome individually. Further annotation as in panel A. (D) Quantification of the combined 4C-seq signal on the short arms of the five acrocentric chromosomes (“intra-arm”), where the rDNA clusters are located, versus the long arms (“inter-arm”) in MLN8237 treated cells and controls at 1.5 h after mitotic release. (E) Zoomed-in normalized and binned intra-chromosomal 4C-seq signal at 500 kb resolution for the rDNA clusters on the five acrocentric chromosomes. The difference between panels is indicated in-between (log2 ratio). Normalization was performed for each chromosome individually. Further annotation as in panel A. (F) Quantification of differences in normalized inter-arm contacts for replicate samples at 1.5 h after mitotic release.

